# Mechanism for controlled assembly of transcriptional condensates by Aire

**DOI:** 10.1101/2023.10.23.563603

**Authors:** Yu-San Huoh, Qianxia Zhang, Ricarda Torner, Sylvan C. Baca, Haribabu Arthanari, Sun Hur

**Affiliations:** Howard Hughes Medical Institute and Program in Cellular and Molecular Medicine, Boston Children’s Hospital, MA 02115, USA; Department of Biological Chemistry and Molecular Pharmacology, Harvard Medical School, Boston, MA 02115, USA; Department of Cancer Biology, Dana-Farber Cancer Institute, Boston, MA 02115, USA; Department of Medical Oncology, Dana-Farber Cancer Institute, Boston, MA 02115, USA

## Abstract

Transcriptional condensates play a crucial role in gene expression and regulation^1–4^, yet the mechanisms governing their assembly at the correct location and time remain poorly understood^2,5–7^. We here report a multi-layered mechanism for condensate assembly by the Autoimmune regulator (Aire), an essential transcriptional regulator (TR) that orchestrates gene expression reprogramming for central T-cell tolerance^8,9^. Previous studies demonstrated that Aire utilizes its polymerizing domain CARD to form nuclear condensates critical for its transcriptional activity^10,11^. Our current data show that Aire condensates assemble on enhancers, stimulating local transcriptional activities, while also connecting disparate inter-chromosomal genomic loci. This process of functional condensate formation hinges upon the coordination between three Aire domains: the CARD, the histone binding domain PHD1 and the activation domain C-terminal tail (CTT). Specifically, CTT directly binds CBP/p300 coactivators, recruiting Aire to CBP/p300-rich enhancers and promoting CARD-mediated condensate assembly. Conversely, PHD1 restrains spontaneous Aire polymerization by binding to the ubiquitous unmethylated H3K4, ensuring Aire remains dispersed throughout the genome until it nucleates on enhancers. Accordingly, deletion of CTT or PHD1 leads to a complete loss of Aire’s functions through distinct mechanisms––deletion of CTT abrogates Aire condensate formation, whereas deletion of PHD1 leads to dysfunctional Aire condensates detached from chromatin. Thus, our findings highlight the balance between PHD1-mediated suppression and CTT-mediated stimulation of Aire polymerization to form transcriptionally active condensates at appropriate target sites, providing new insights into controlled polymerization of TRs.

## Main

Controlled protein polymerization underlies a diverse range of biological processes^12–14^. One family of protein domains that mediate polymerization is the <u>C</u>aspase <u>A</u>ctivation <u>R</u>ecruitment <u>D</u>omain (CARD), which plays important roles in cell death and inflammatory signaling^15,16^. CARD domains often self-polymerize into filaments, serving as a key mechanism to amplify upstream immune signals^15^. However, aberrant polymerization of CARD can lead to undesirable consequences such as chronic inflammation or toxicity^17–19^. Thus, tight regulatory mechanisms are required to ensure that CARD polymerization occurs only under specific conditions^15,17,18^.

Despite their well-known roles in cytosolic signaling pathways, CARDs are understudied in transcriptional regulators (TRs) like Aire and Speckle Proteins (SPs)^10,20^. Consequently, specific functions and mechanisms regulating CARD polymerization in transcription remain poorly understood.

Aire plays a critical role in central T cell tolerance^8,9^. Aire orchestrates the expression of thousands of peripheral tissue antigens (PTAs) in medullary thymic epithelial cells (mTECs)^21,22^. These PTAs are displayed on the mTEC cell surface to recognize auto-reactive T cells for their negative selection or diversion into regulatory T cells^8,21,23,24^. Consequently, mutations in human *AIRE* or knock-out of mouse *Aire* result in multi-organ autoimmunity, including autoimmune polyendocrinopathy syndrome type 1 (APS-1)^21,25^. Initially, Aire was regarded as a conventional transcription factor (TF) directly binding PTA gene promoters to induce expression^26,27^. Recent studies, however, propose that Aire largely upregulates PTA expression indirectly by amplifying the actions of various lineage-defining TFs ectopically expressed in mTECs^28–32^. Precise modes of action and the underlying mechanisms for Aire’s synergy with these TFs remain unclear.

Aire is a chromatin-binding TR, rather than a sequence-specific, direct DNA binder^33,34^. Although Aire possesses a putative DNA binding domain, SAND, this domain lacks essential DNA binding residues^35^. Instead, Aire features two potential chromatin reader domains––PHD1 and PHD2. PHD1 binds histone H3 (H3) with non-methylated lysine 4 (H3K4me0), a histone state that is depleted from active loci but is abundantly present elsewhere^36–38^. Conversely, PHD2 exhibits little histone binding activity, and its functions remain unknown^33,34,39^. At first, PHD1’s specificity for H3K4me0 was thought to guide Aire to inactive PTA gene loci^40,41^. However, a study showed that Aire primarily binds genomic sites pre-enriched with the permissive histone mark, H3K27ac, notably at super-enhancers (SEs)^42^. Confoundingly, regions rich in active H3K27ac marks tend to lack H3K4me0^43,44^, raising questions about how Aire specifically targets H3K27ac-rich sites and the precise role of PHD1 in this context.

One of the most intriguing properties of Aire as a TR is its ability to form nuclear condensates. These condensates are easily visualized by diffraction-limited light microscopy in both human and murine mTECs as well as in model cell lines^10,11,45–47^. A previous study showed that Aire forms homopolymers using its N-terminal CARD^11^. This homopolymerization closely correlates with Aire nuclear condensate formation^11^, suggesting Aire homopolymers manifest as nuclear condensates. Additionally, Aire CARD can be functionally substituted with an orthogonal, chemically-inducible multimerizing domain, preserving both the condensate formation and transcriptional activity^11^. Notably, substitution with a mere dimerization or tetramerization domain is insufficient^11^. These observations, in combination with other reports^10,48,49^ demonstrate the importance of Aire polymerization in condensate formation and transcriptional activity. However, while Aire condensate formation is necessary, it is not sufficient for transcriptional function. Equally critical appears to be the localization of Aire condensates, as Aire condensates associated with PML bodies leads to the loss of Aire transcriptional activity^11^. In fact, the precise locations and functions of Aire condensates remain unclear; it is debated whether Aire condensates form at Aire-bound genomic loci and whether these condensates serve as active transcription sites, inactive storage depots, or suppressive compartments^42,50,51^.

In this study, we demonstrate that Aire condensates indeed assemble on enhancers, serving as hubs for transcriptional activation. Moreover, we reveal that these condensates are subject to intricate regulatory mechanisms ensuring tight coordination of CARD polymerization with genomic target recognition.

### Aire condensates form on enhancers and activate transcription

Aire is known to be expressed in a miniscule subset of mTECs with temporal dynamics^29–31,52,53^, which has made mechanistic studies using mTECs challenging. Therefore, we generated a doxycycline (Dox)-inducible model system where Aire is ectopically expressed in human thymic epithelial cell line (4D6) at levels equivalent to that in human mTECs (Fig. 1a). It is worth noting that the mRNA level of endogenous Aire in mTECs matches that of highly abundant proteins, such as ribosomal proteins, actin and GAPDH, which we reproduced in our 4D6 cells. Our 4D6 system also recapitulated Aire localization to H3K27ac-enriched sites including SEs, as measured by Aire and H3K27ac ChIP-seq (Fig. 1b and Extended Data Fig. 1a)^42^. It also showed CARD-dependent nuclear condensate formation (Fig. 1e), Aire-induced broad transcriptomic changes (Extended Data Fig. 1b) and the impact of loss-of-function APS-1 mutations (Extended Data Fig. 1c)––characteristics that were all previously observed in mTECs^26,42,47,53–56^. We thus utilized 4D6 cells to investigate the mechanism of Aire polymerization in the nucleus.

**Fig. 1.**
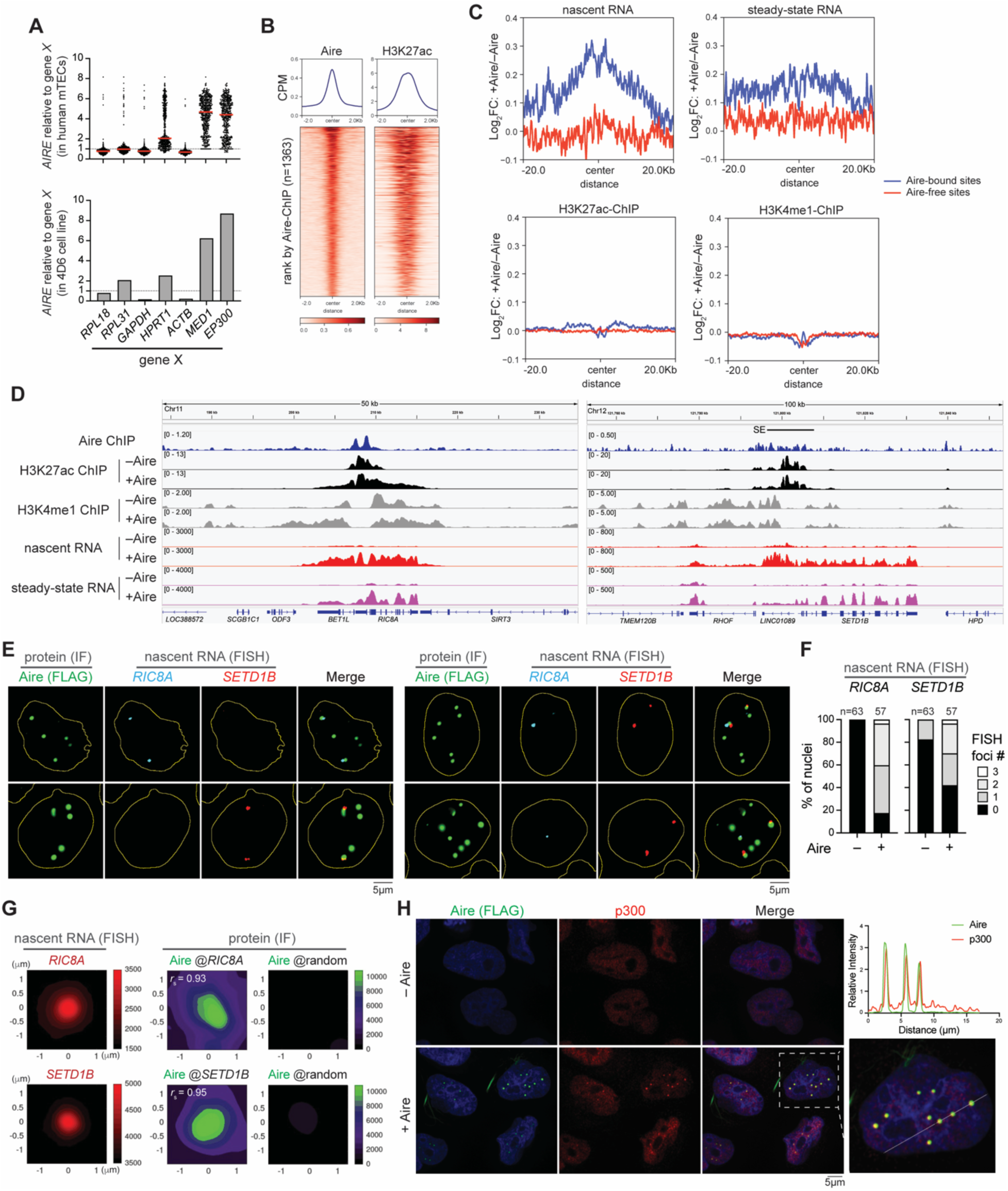
Aire condensates form on enhancers and activate transcription. (A) *AIRE* transcripts relative to other gene (X) transcripts in human *AIRE*^+^ mTECs (top panel) and Dox-inducible *AIRE*-expressing 4D6 cells (bottom panel). Bulk RNA-seq was performed on 4D6 cells 24 hrs post-induction of *AIRE* expression and compared to previously published single-cell RNA-seq data of human thymic epithelial cells (n = 477 *AIRE*^+^ mTECs)^1^. (B) Heatmaps of normalized Aire and H3K27ac ChIP-seq signals (Counts Per Million, CPM) in 4D6 cells. Heatmaps are centered on all Aire peaks (n = 1363) and ranked by Aire ChIP-seq intensity. Aire ChIP-seq was performed on 4D6 cell line expressing WT human Aire-FLAG (Aire-FLAG) under a doxycycline (Dox)-inducible promoter. H3K27ac ChIP-seq was performed on the same 4D6 cells prior to Aire expression (pre-Aire). Unless mentioned otherwise, Aire indicates human Aire throughout the manuscript. (C) Aire-induced changes in transcription or histone marks 20 kb upstream or downstream of Aire-bound and Aire-free nucleosome-free regions (NFRs, n = 542 and 658, respectively). FC, fold-change. (D) Genome browser views of normalized ChIP-seq and RNA-seq profiles at exemplar Aire-bound sites in 4D6 cells. Numbers to the left indicate the ranges of normalized reads for RNA-seq or counts per million (CPM) for ChIP-seq. SE, super-enhancer. (E) Nascent RNA-fluorescence in situ hybridization (FISH) coupled with immunofluorescence (IF) images of stable 4D6 cells expressing Aire-FLAG under a Dox-inducible promoter. Cells were induced with 1 µg/ml Dox (+Aire) for 24 hrs before staining with anti-FLAG along with FISH probes targeting *RIC8A* and *SETD1B*. RNA-FISH probes were designed to hybridize with the intronic regions of Aire-dependent targets (Supplementary table 2). Yellow outlines represent boundaries of nuclei determined by DAPI staining. Representative images highlight nuclei containing Aire condensates (IF) with either *RIC8A* (top left panels) or *SETD1B* (bottom left panels) RNA-FISH foci, or nuclei containing Aire condensates (IF) along with both RNA-FISH foci (top and bottom right panels). See Extended Data Fig. 2A for staining of cells that were not induced with Dox (–Aire) in addition to a zoomed-out image of +Aire cells. (F) Quantitation of percent of nuclei that have 0, 1, 2 or 3 RNA-FISH foci in 4D6 cells before and after Aire expression as shown in (E) and Extended Data Fig. 2A. n indicates the number of nuclei examined. (G) Spatial relationship between RNA-FISH foci and Aire condensates. Shown are average signals of RNA-FISH (left), Aire IF centered on the indicated FISH foci (center) and Aire IF centered on randomly selected nuclear positions (right). *r*_s_ denotes the Spearman’s correlation coefficient between RNA-FISH and Aire IF signals. (H) Representative immunofluorescence images of endogenous p300 in Dox-inducible stable 4D6 cells. Cells were not (– Aire) or were induced with 1 µg/ml Dox (+ Aire) for 24 hrs before immunostaining with anti-FLAG and anti-p300. Right: zoomed-in view of a nucleus enclosed by a white-dashed box along with measured fluorescence intensities across a drawn solid white line. See Extended Data Fig. 3A for IF images of endogenous CBP and MED1 the same 4D6 cell line. All data are representative of at least three independent experiments.

To characterize Aire’s molecular functions, we first examined the impact of Aire at Aire-bound genomic regions by assessing the changes in both the steady state (by bulk RNA-seq) and nascent (by 5-ethynyl uridine (5EU)-seq) RNA levels upon Aire expression. Aire-bound vs. Aire-free regions were defined as nucleosome-free regions (by ATAC-seq) that were highly occupied by Aire vs. regions lacking Aire ChIP-seq signals (Supplementary Table 1). These Aire-bound and Aire-free regions were chosen to ensure comparable chromatin accessibility (Extended Data Fig. 1d). There was a global increase in both nascent and steady-state RNAs at the Aire-bound loci upon Aire expression, but not in Aire-free regions (Fig. 1c, top two panels). Notably, the Aire-mediated transcriptional induction was more pronounced and more focused around the Aire-bound site, when examining the nascent RNAs than the steady-state RNAs. Given that Aire primarily localizes at active enhancers (Fig. 1b and Extended Data Fig. 1a), this suggests that Aire-mediated transcriptional activation occurs largely in the form of short-lived enhancer RNAs (eRNAs). Analyses of several loci, such as *RIC8A, SETD1B*, *UBTF*, and *STX10*, which were all up-regulated and bound by Aire, also showed marked increase in nascent transcripts upon Aire expression (Fig. 1d and Extended Data Fig. 1f). Intriguingly, this Aire-dependent transcriptional activation at Aire-bound loci was not always accompanied by an increase in H3K4me1 or H3K27ac level as assessed by histone mark ChIP-seq. At the *RIC8A* locus, Aire induced spreading of H3K4me1 and H3K27ac, whereas at *SETD1B*, *UBTF* and *STX10* loci, no such changes were observed (Fig. 1d and Extended Data Fig. 1f). Global analysis of H3K4me1, H3K27ac and H3K4me3 ChIP and ATAC signals aggregated over Aire-bound sites also showed minimal changes in these histone marks or chromatin accessibility upon Aire occupancy (Fig. 1c and Extended Data Fig. 1e).

To examine whether the transcriptional activation of Aire-bound genomic regions occurs within Aire condensates, we performed immunofluorescence (IF) combined with nascent RNA fluorescence in situ hybridization (FISH) to examine *RIC8A, SETD1B*, and *UBTF* loci (Supplementary Table 2). In ∼80% of the cells examined, Aire expression resulted in the appearance of 1 or 2 nascent RNA-FISH foci for *RIC8A, SETD1B* and *UBTF* (Fig. 1e,f and Extended Data Fig. 2a,b). This is consistent with the transcriptional activation of these genes by Aire as measured with 5EU-seq and total RNA-seq (Fig. 1d and Extended Data Fig. 1f). RNA-FISH foci and Aire condensates had significant overlap as demonstrated by the averaged Aire fluorescence intensities centered on RNA-FISH foci (Fig. 1g and Extended Data Fig. 2c) and individual distance between the centers of the closest RNA-FISH foci and Aire condensate (Extended Data Fig. 2d). Additionally, Aire condensates also colocalized with coactivators p300, CBP and MED1 (Fig. 1h and Extended Data Fig. 3a). Similar condensates of p300 and CBP were not detected in the absence of Aire, although MED1 condensates were seen even without Aire (Fig. 1h and Extended Data Fig. 3a). These results together suggest that Aire condensates are indeed composed of genomic sites bound and activated by Aire.

Among all Aire-positive nuclei, 49% showed both *RIC8A* FISH foci and *SETD1B* FISH foci within the same nucleus (Fig. 1e, right panels); within these nuclei containing both *RIC8A* and *SETD1B* FISH foci, ∼20% of the *RIC8A* and *SETD1B* FISH foci shared an Aire condensate with each other (Extended Data Fig. 2e). We observed a similar frequency of sharing the same Aire condensate between *RIC8A* and *UBTF* FISH foci (Extended Data Fig. 2f). While these observations do not support obligatory pairings among these genes, given that *RIC8A, SETD1B* and *UBTF* are located on different chromosomes, their frequent contacts through the same Aire condensates suggests that each cluster of Aire is likely connecting distinct inter-chromosomal genomic loci to form a transcriptional condensate.

### Aire condensate formation requires the activation domain C-terminal tail (CTT)

We next examined the mechanism by which Aire forms condensates. Systematic domain truncation analysis showed that CARD was indispensable for Aire condensate formation (consistent with previous reports^10,11^), whereas SAND, PHD1 and PHD2 were dispensable (Fig. 2a,b and Extended Data Fig. 3b). Intriguingly, deletion of C-terminal tail (CTT, residues 482-545 in human and 480-552 in mouse) completely abrogated condensate formation for both human and mouse Aire (Fig. 2b and Extended Data Fig. 3b). While the Aire CARD domain spontaneously polymerizes *in vitro*^11^, isolated CTT behaves as a monomer. Solution NMR of isolated CTT showed a 15N T_2_ relaxation time (Extended Data Fig. 3c) comparable to other monomeric activation domains of similar size^57,58^. Furthermore, unlike isolated CARD that forms nuclear condensates^11^, isolated CTT (tagged with an artificial protein, APEX2-GST) did not show condensates in 293T cells (Extended Data Fig. 3d). These observations indicate that CTT may instead modulate CARD polymerization, rather than directly participate in Aire condensate formation.

**Fig. 2.**
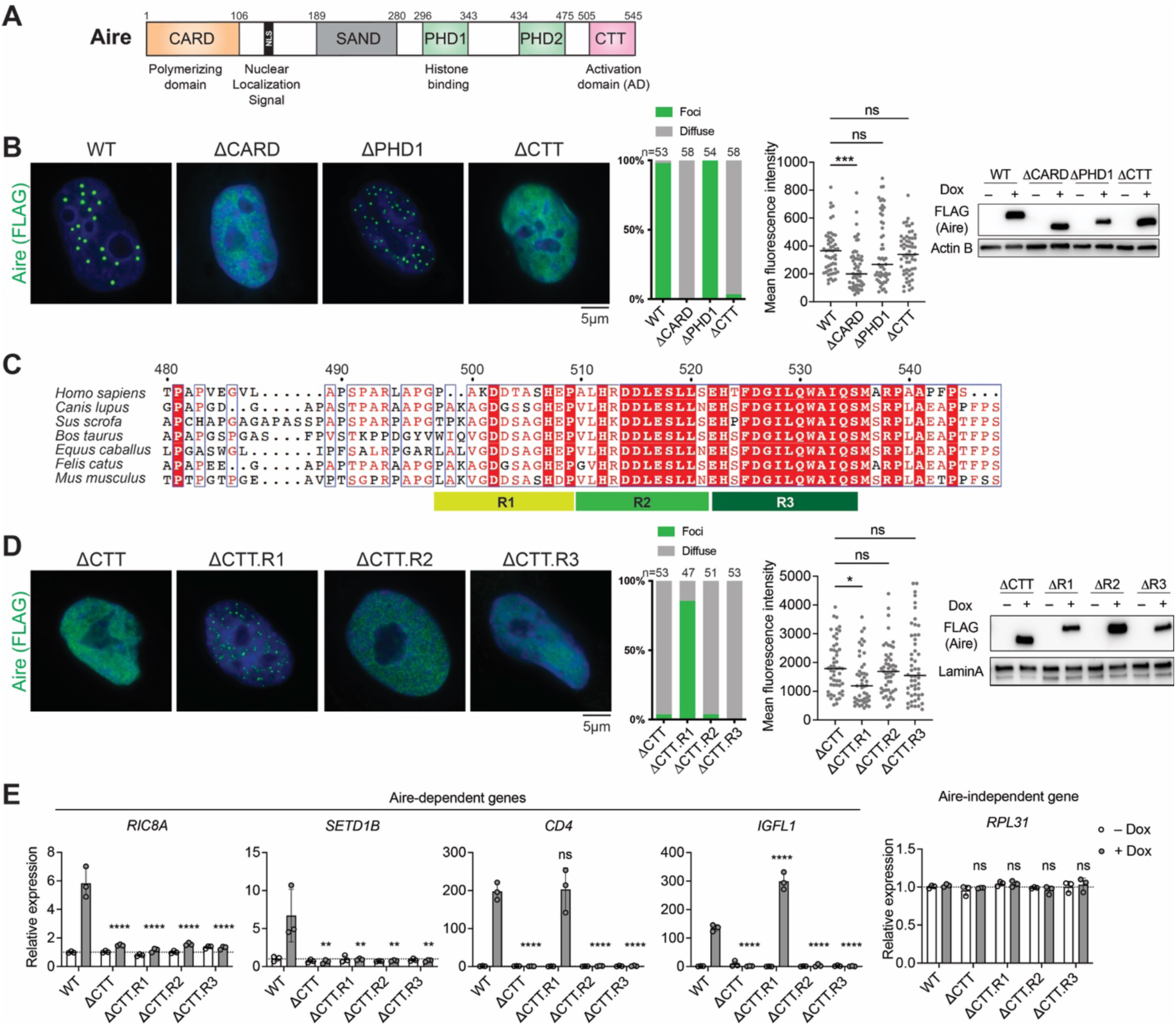
Aire condensate formation requires the AD-like C-terminal tail (CTT). (A) Schematic of Aire domain architecture with previously characterized functions of individual domains. The numbers above denote amino acid residuals of human Aire (Aire). (B) Representative immunofluorescence images of Aire ΔCARD, ΔPHD1 and ΔCTT variants in Dox-inducible stable 4D6 cells. Middle: percentage of nuclei with Aire condensates vs. diffuse Aire staining; and mean fluorescence intensity of nuclei examined. n represents the number of nuclei examined for each sample. *p*-values (Kruskal-Wallis test with Dunn’s multiple comparisons test) were calculated in comparison to WT Aire. \*\*\**p* < 0.001; *p* > 0.05 is not significant (ns). Right: Western blot (WB) showing the levels of FLAG-tagged Aire variants. (C) Sequence alignment of Aire CTT domains from various species. CTT.R1, R2 and R3 indicate aa 499-509, aa 510-521, aa 522-535, respectively, in Aire. (D) Representative immunofluorescence images of Aire 1′CTT, ΔCTT.R1, ΔCTT.R2, and ΔCTT.R3 in 4D6 cells. Middle: percentage of nuclei with Aire condensates vs. diffuse Aire staining; and mean fluorescence intensity of nuclei examined. n represents the number of nuclei examined for each sample. *p*-values (Kruskal-Wallis test with Dunn’s multiple comparisons test) were calculated in comparison to AireDCTT. \**p* < 0.05; *p* > 0.05 is not significant (ns). Right: WB showing the nuclear expression levels of FLAG-tagged Aire variants. Experiments were done as in (B). (E) Transcriptional activity of WT Aire or various CTT deletion mutants, as measured by the relative mRNA levels of Aire-dependent genes, *RIC8A, SETD1B*, *CD4,* and *IGFL1* in Dox-inducible stable 4D6 cells. An Aire-independent gene, *RPL31*, was examined as a negative control. All genes were normalized against the internal control *RPL18*. Data are presented as mean ± SD, n = 3. *p*-values (one-way ANOVA with Dunnett’s multiple comparisons test) were calculated in comparison to WT Aire \*\**p* < 0.01; \*\*\*\**p* < 0.0001; *p* > 0.05 is not significant (ns). All data are representative of at least three independent experiments.

To determine which region within CTT is important for Aire condensate formation, we generated 4D6 cell lines expressing Aire CTT truncation mutants under a Dox-inducible promoter, as was done for WT Aire and other domain deletion mutants. Further truncation analysis suggests that residues 510-521 (R2, human residue numbering) and 522-535 (R3) within CTT were required for Aire condensate formation, but residues 499-509 (R1) of CTT were not (Fig. 2c,d). R2 and R3 also displayed higher sequence conservation than R1 and the rest of the CTT (Fig. 2c), suggesting that R2-R3 may have different functions than R1.

Aire CTT is known to have transcriptional activation domain (AD)-like activity^59,60^. We thus asked how the same CTT truncations affect AD-like activity. We measured the CTT variants’ transcriptional activities by RT-qPCR of several Aire target genes. Deletion of R2 or R3 (CTTΔR2 and ΔR3) completely abolished Aire’s transcriptional activity regardless of the examined target genes (Fig. 2e), recapitulating the loss-of-function phenotype observed with the complete deletion of CTT (ΔCTT). On the other hand, CTTΔR1 showed gene-specific behaviors, suggesting a more nuanced function for R1 (Fig. 2e). An AD reporter assay using CTT-fused with Gal4 DNA-binding domain (Gal4^DBD^)^59^ also highlighted the importance of R2 and R3 in CTT’s AD-like activity (Extended Data Fig. 3e), although R1 was also important in this reporter assay.

Collectively, these results suggest distinct functions for CTT R1 and R2-R3. R2-R3 contribute to both condensate formation and the AD-like activity, whereas CTT R1 is involved in transcriptional activation of a subset of target genes, with minimal impact on condensate formation (Fig. 2d,e and Extended Data Fig. 3e).

### Aire CTT directly binds CBP/p300 for transcriptional activity and condensate formation

To elucidate how CTT promotes Aire condensate formation and transcriptional activity, we investigated both the genetic and physical interaction partners of CTT. Based on our Gal4-based Aire CTT reporter assay (Extended Data Fig. 3e), we designed a genome-wide CRISPR screen. This screen utilized a 4D6 cell line with the stable incorporation of the fluorescent reporter mKate2 under the control of upstream activation sequences (UAS). The mKate2 reporter was induced upon the expression of Gal4^DBD^-CTT fused with the expression reporter GFP via a self-cleavable peptide 2A (Fig. 3a). We then collected GFP-positive cells that had decreased or increased mKate2 expression after transducing lentiviral sgRNA libraries and compared sgRNA enrichment between these two populations (Fig. 3b). In parallel, we looked for physical Aire CTT interaction partners by performing GST pull-downs of 293T lysate using purified GST-tagged CTT (GST-CTT) and analyzed the co-purified proteins by mass-spectrometry (Fig. 3c). From these two independent analyses, we identified the transcriptional co-activators CBP and p300 as the most significant common hits (Fig. 3b,c and Extended Data Fig. 4a; Supplementary Tables 3,4). CBP and p300 are highly homologous histone acetyltransferases (HATs) responsible for producing a large portion of H3K27ac in cells^61^. CBP/p300 colocalize with Aire condensates in both mTECs^10,46^ and 4D6 cell line (Fig. 1h and Extended Data Fig. 3a). No other HATs were found in either screen.

**Fig. 3.**
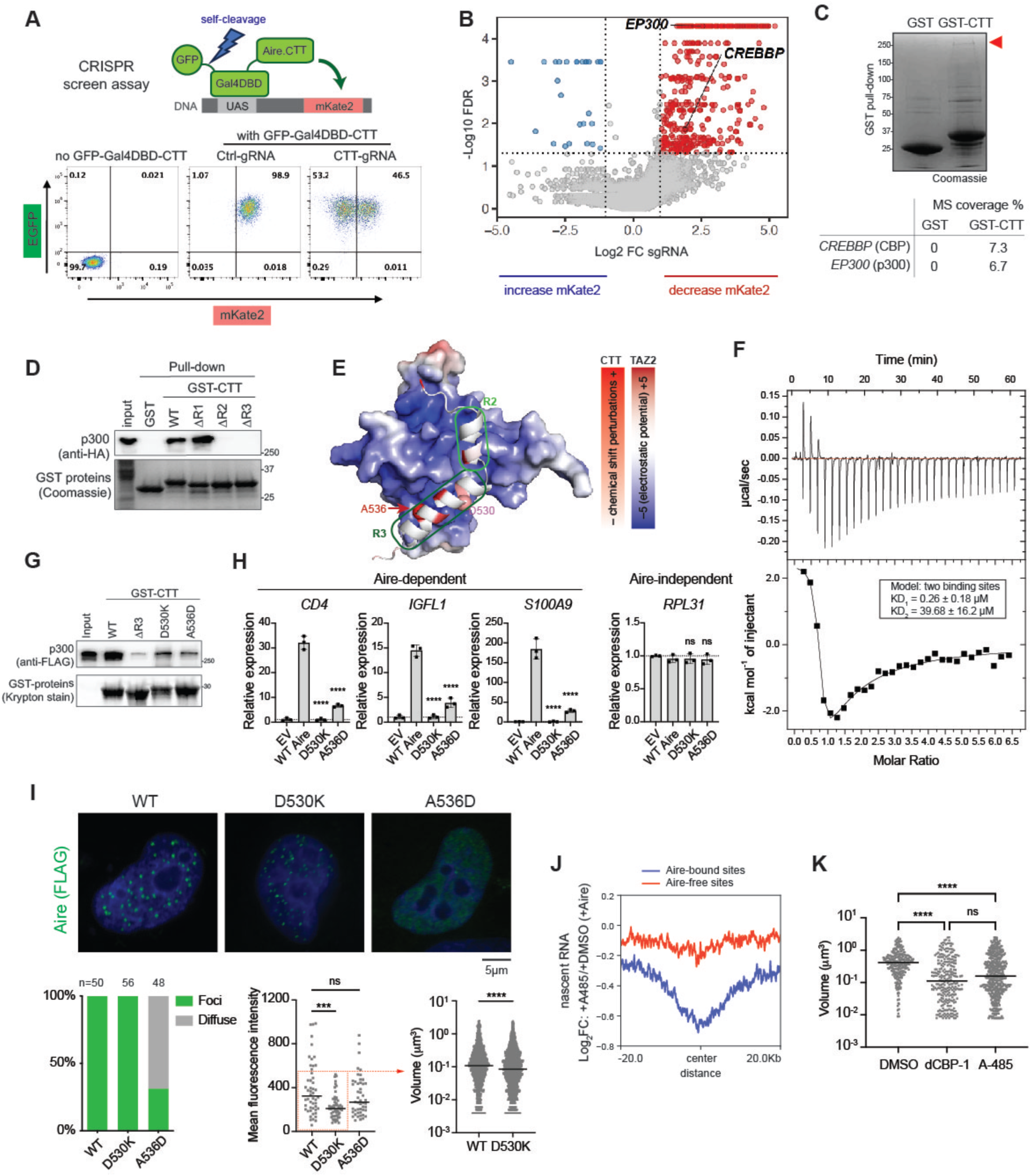
Aire CTT directly binds CBP/p300 for transcriptional activity and condensate formation. (A) Gal4-reporter based CRISPR screening assay in 4D6 cells. Gal4-DNA binding domain (Gal4DBD)-CTT from Extended Data Fig. 3D was fused with GFP via a self-cleavable peptide (P2A). Dox-inducible expression of GFP-Gal4DBD-CTT leads to the expression of mKate2 under the control of UAS. Bottom: validation of the system by flow cytometry. Ctrl, control; gRNA, guide RNA. (B) Single gRNA (sgRNA) enrichment in 4D6 cells with decreased (red) versus increased (blue) expression of mKate2. FDR, false discovery rate. FC, fold-change. See Supplementary Table 3 for the comprehensive list of sgRNA enrichment results. (C) Coomassie Blue-stained SDS-PAGE gel of GST pull-down eluates. Purified recombinant His_6_-GST-CTT (or His_6_-GST) was incubated with 293T nuclear extracts prior to pull-down. Red arrow denotes the location of the ∼250 kDa gel band that was cut and analyzed by mass spectrometry. Bottom: sequence coverages determined by mass spectrometry (MS) of p300 and CBP pulled-down with GST or GST-CTT. See Extended Data Fig. 4A and Supplementary Table 4 for the comprehensive list of top hits. (D) His_6_-GST and His_6_-GST-CTT variant pull-downs of HA-tagged p300 transiently expressed in 293T nuclear extracts. (E) NMR results incorporated into the top AlphaFold model^2^ of the CBP–CTT complex using CBP TAZ2 (aa 1764-1855) and mouse Aire CTT (aa 480-552). CTT is colored based on the degree of NMR chemical shift perturbation upon TAZ2 interaction. CBP TAZ2 is colored by electrostatic potential (PyMol, APBS module). Mouse Aire CTT residues D530 and A536 within R3 are highlighted with spheres. (F) Isothermal titration calorimetry thermograms of Aire CTT titrated into CBP TAZ2 and fit to a two-sites binding model. (G) His_6_-GST-CTT variant pull-downs of full-length FLAG-tagged p300. Both baits and prey were purified recombinant proteins. (H) Transcriptional activity of WT Aire or CTT point mutants, as measured by the relative mRNA levels of Aire-dependent genes, *CD4*, *IGFL1* and *S100A9*, in 4D6 cells. Cells were transiently transfected with plasmids expressing mouse Aire and its variants. An Aire-independent gene, *RPL31*, was also examined as a negative control. All genes were normalized against the internal control *RPL18*. Data are presented as mean ± SD, n = 3. *p*-values (one-way ANOVA with Dunnett’s multiple comparisons test) were calculated in comparison to WT Aire. \*\*\*\**p* < 0.0001; *p* > 0.05 is not significant (ns). (I) Representative immunofluorescence images of FLAG-tagged Aire variants in 4D6 cells. Cells were transfected with mouse Aire-FLAG expression plasmids 24 hrs prior to fixation. Bottom left: percentage of nuclei with Aire condensates vs. diffuse Aire staining. Bottom middle: mean fluorescence intensity of all Aire-expressing nuclei examined. Nuclei within the red box (fluorescence intensity ≤550 a.u.) were selected to ensure that condensate size comparisons were from cells with similar Aire expression levels. Note that the fluorescent intensities between the selected WT and D530K nuclei were not significantly different. Bottom right: quantification of Aire condensate volumes in nuclei with similar mean fluorescent intensities. See Extended Data Fig. 6B for quantification of Aire condensates volumes in all nuclei examined. *p*-values (Kruskal-Wallis test with Dunn’s multiple comparisons test for the bottom-middle panel and Mann-Whitney test for the bottom-right panel) were calculated in comparison to WT Aire. \*\*\**p* < 0.001; \*\*\*\**p* < 0.0001; *p* > 0.05 is not significant (ns). (J) Aire-induced transcriptional changes in the presence or absence of A-485 20 kb upstream or downstream of Aire-bound or Aire-free nucleosome-free regions (NFRs). Dox, DMSO or A-485 (3 µM), and 5’-EU were added to cell culture 24 hrs, 4hrs, and 0.5 hr prior to RNA extraction, respectively. (K) Quantification of Aire condensates volumes (bottom left) in the presence of the p300/CBP degrader dCBP-1 (0.25 µM) or the catalytic inhibitor A-485 (3 µM) in comparison to vehicle control DMSO. Dox was added to induce expression of Aire for 8 hrs prior to fixation, and DMSO, dCBP-1, or A-485 were added 4 hrs prior to fixation. Aire condensates n = 249, 259, 502 in DMSO–, dCBP-1– and A-485–treated cells, respectively. *p*-values (Kruskal-Wallis test with Dunn’s multiple comparisons test) were calculated. \*\*\*\**p* < 0.0001; *p* > 0.05 is not significant (ns). See Extended Data Fig. 6D for Representative immunofluorescence images of WT Aire-FLAG in Dox-inducible 4D6 cells.

In comparison to intact CTT, CTTΔR2 and ΔR3 showed reduced CTT–CBP/p300 interaction, whereas CTTΔR1 maintained similar binding levels (Fig. 3d). Thus, unlike R1, the importance of R2 and R3 in CBP/p300 binding correlated with their significance in Aire condensate formation (Fig. 2d). Using recombinant CTT and p300, we verified that the interaction was direct (Fig. 3e-g). Domain truncation analysis of p300 revealed that CTT interacted with the TAZ2 and IBiD domains of p300 (Extended Data Fig. 4b,c). Isolated CBP TAZ2 could be recombinantly purified and showed direct binding to purified CTT (Extended Data Fig. 4d), while isolated IBiD was insoluble, precluding more detailed binding analysis. Isothermal Titration Calorimetry (ITC) analysis showed that Aire CTT binds CBP TAZ2 with a *K*_d_ of 0.26 mM (Fig. 3f), consistent with the TAZ2 affinity of other ADs^62,63^. Note that ITC also detected a second, low-affinity binding site (of 40 mM, Fig. 3f), which is presumably not as important as the high-affinity site.

To further characterize the TAZ2–CTT interaction, we performed ^1^H-^15^N HSQC NMR spectroscopy on CTT with and without TAZ2. In the absence of TAZ2, isolated CTT displayed largely disordered characteristics (Extended Data Fig. 5a), although R2 and R3 are predicted to have a moderate tendency to form alpha helices (Fig. 3e and Extended Data Fig. 5b). Upon incubation with CBP TAZ2, R2 and R3 underwent significant chemical shifts and peak broadening (Extended Data Fig. 5c,d), suggesting that R2 and R3 bind TAZ2. The utilization of R2 and R3 for TAZ2 binding was further supported by AlphaFold prediction (Fig. 3e).

AlphaFold model also predicted alpha helical conformations of R2 and R3, akin to the binding mode observed for other ADs^64,65^. Furthermore, mutations in the putative CTT interface (mouse Aire D530K and A536D, corresponding to human Aire D526 and A532) significantly reduced the CTT–CBP interaction, as measured by pull-down assay using purified proteins (Fig. 3g). This mutational analysis provides additional support for the NMR data and AlphaFold model (Fig. 3e).

We next examined whether CBP/p300 play an important role in Aire transcriptional activity and condensate formation. Transient expression of WT vs. D530K or A536D showed that the mutations significantly lowered the transcriptional activities of Aire (Fig. 3h) and the AD reporter activity of Gal4^DBD^-CTT (Extended Data Fig. 6a). IF analysis also showed that A536D was diffuse in most cells (Fig. 3i). D530K formed condensates at a similar frequency as WT Aire, but D530K condensates were smaller than WT condensates, irrespective of whether cells had similar Aire expression levels (as defined by similar nuclear intensities, Fig. 3i) or not (Extended Data Fig. 6b). These results show that both A536D and D530K are impaired in Aire condensate formation, albeit to differing degrees.

To further test the role of CBP/p300 in Aire functions, we used two pharmacological inhibitors of CBP/p300, the catalytic inhibitor A-485^66^ and the degrader dCBP-1^67^. As expected, treatment with A-485 for 4 hrs reduced the levels of H3K27ac without diminishing the CBP/p300 protein levels, whereas dCBP-1 significantly reduced the levels of both H3K27ac and CBP/p300 (Extended Data Fig. 6d). Global analysis of nascent RNAs (by 5EU-seq) showed that A-485 significantly reduced transcriptional activity at Aire-bound genomic regions, but not at Aire-free regions with comparable chromatin accessibilities (Fig. 3j and Extended Data Fig. 1d). 5EU-qPCR of Aire target genes showed that dCBP-1 also had a similar negative impact on transcription of Aire-bound loci (Extended Data Fig. 6c). IF of Aire with and without CBP/p300 inhibitors showed that both dCBP-1 and A-485 treatment significantly decreased the size of Aire condensates, all without affecting Aire expression level (Extended Data Fig. 6d). Furthermore, the smaller condensates visible with dCBP-1 and A-485 were transcriptionally inactive, as evidenced by the lack of nascent RNA-FISH foci of an Aire target gene, *RIC8A* (Extended Data Fig. 6e). Similarly, dCBP-1 and A-485 decreased the AD reporter activity of Gal4^DBD^-CTT, despite slightly increasing the level of Gal4^DBD^-CTT (Extended Data Fig. 6f,g). This inhibition was specific to CBP/p300, as inhibition of Brd4, which often cooperates with CBP/p300 for enhancer functions^68^, did not inhibit Aire’s transcriptional activity or condensate formation (Extended Data Fig. 6h,i), consistent with a previous report^69^.

Overall, these results suggest that Aire CTT’s role in Aire condensate formation and transcriptional activity are mediated by CBP/p300.

### CTT directs Aire to enhancers pre-enriched with CBP/p300

We next aimed to elucidate how the CTT–CBP/p300 interaction facilitates both Aire condensate formation and transcriptional functions. Our attention turned to Aire’s ability to bind H3K27ac-rich enhancer loci, which are regions also known to have a high occupancy of CBP/p300. We hypothesized that the clustering of CBP/p300 might play a role in recruiting Aire to these enhancer loci and could nucleate Aire polymerization by increasing the local concentration of Aire molecules. To investigate the role of CBP/p300 in Aire’s enhancer targeting, we performed p300 ChIP-seq using Aire-inducible 4D6 cells prior to Aire expression in order to compare the “pre-Aire” distribution of p300 with Aire distribution patterns. The result revealed a significant enrichment of p300 at Aire ChIP peaks, even before Aire expression (Fig. 4a). When we compared Aire ChIP-seq signals with CBP/p300 or H3K27ac ChIP signals, we observed a closer correlation between Aire binding and pre-existing CBP/p300 occupancy than with H3K27ac occupancy (Fig. 4b, left panels). The discrepancy between CBP/p300 and H3K27ac can be rationalized by the fact that H3K27ac can be generated by other HATs besides CBP/p300^61,70^, and that not all genomic occupancy of CBP/p300 leads to histone acetylation^71^. These observations suggest that CBP/p300 likely play a role in Aire’s genomic targeting.

**Fig. 4.**
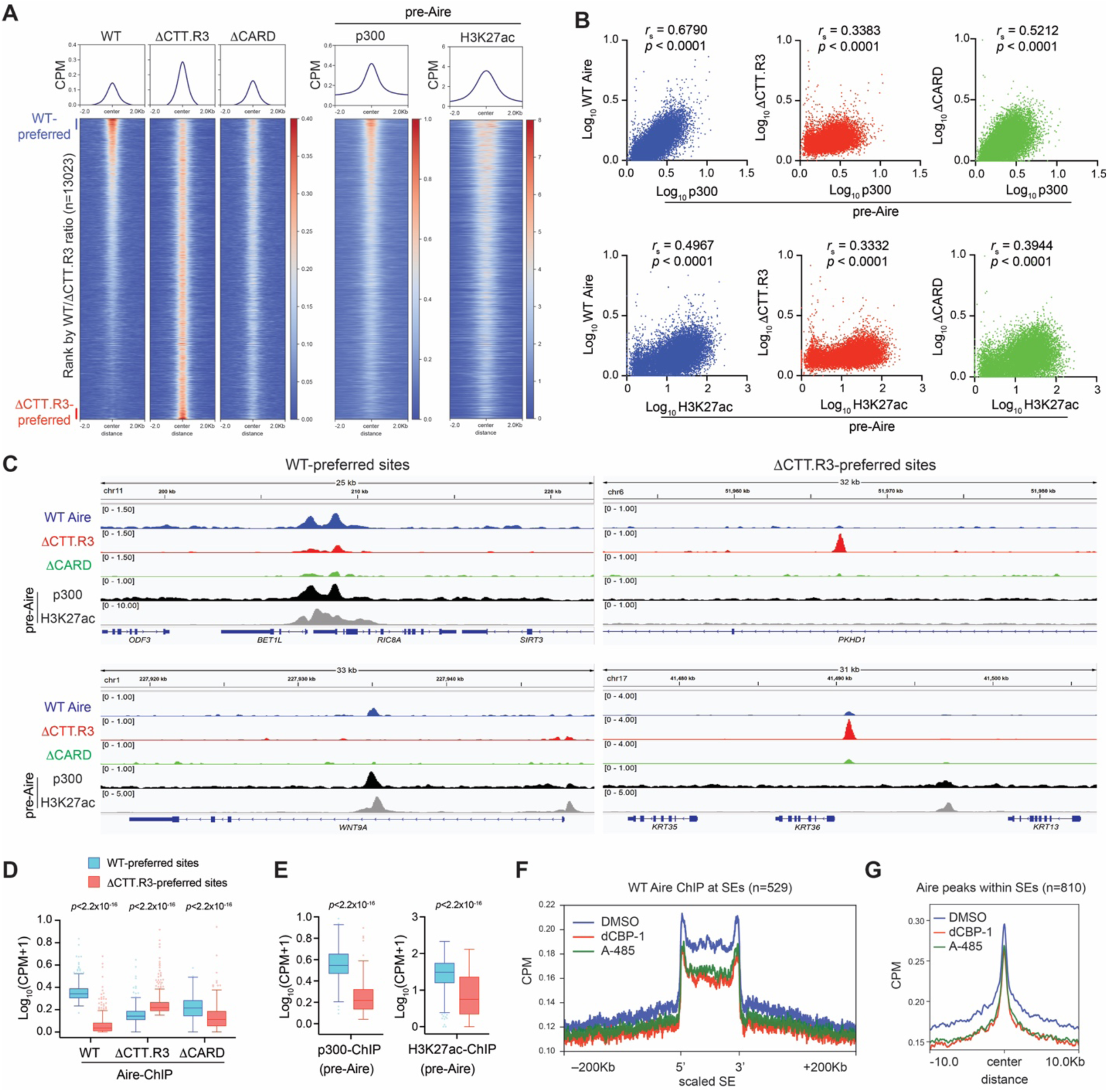
CTT and CARD domains bias Aire towards CBP/p300-enriched genomic loci. (A) Heatmaps of normalized ChIP-seq signals (Counts Per Million, CPM) for indicated proteins in Dox-inducible stable 4D6 cells. Heatmaps are centered on all Aire peaks (n = 13023) and ranked by the ratio of WT Aire over ΔCTT.R3 ChIP-seq signals. Aire ChIP-seq signals were subtracted of background noise from corresponding input controls. p300 and H3K27ac ChIP-seq signals are from 4D6 cells prior to Aire expression (pre-Aire). (B) Correlation between WT Aire, ΔCTT.R3, or ΔCARD with p300 (top panels) or H3K27ac (bottom panels) ChIP-seq signals at all Aire peaks (n=13,023). *r*_s_, Spearman’s correlation coefficient. (C) Genome browser views of normalized ChIP-seq profiles for indicated proteins at exemplar WT-preferred sites versus ΔCTT.R3-preferred sites in Dox-inducible 4D6 cells. Aire ChIP-seq signals shown were subtracted of background noise from corresponding input controls. Numbers to the left of each panel indicate the ranges of normalized CPM for ChIP-seq. (D-E) Quantification of normalized ChIP-seq signals (CPM) for indicated proteins at WT-preferred sites vs. ΔCTT.R3-preferred sites (top and bottom 500 peaks from (A), respectively). ChIP-seq signals were normalized using Trimmed Mean of M-values (TMM, see Methods). *p*-values were calculated using Wilcoxon rank sum test. (F-G) Contribution of CBP/p300 on Aire localization at super-enhancers (SEs). 4D6 cells were treated with Dox for 24 hrs, and DMSO, 0.25 µM dCBP-1 or 3 µM A-485 were added 4 hrs prior to harvest. Average Aire ChIP-seq profiles (normalized CPM) from cells treated with p300 inhibitors compared to DMSO spanning 200Kb up- or downstream of H3K27ac-delimited SEs (n = 529, defined in Extended Data Fig. 1A) (F) or centered at Aire peaks located within SEs (n=810) (G). *p*-values were calculated using Wilcoxon rank sum test where *p* < 2.2×10^-16^ for DMSO vs. dCBP-1 and DMSO vs. A-485 in (F-G).

To further investigate whether CBP/p300 guide Aire’s target site selection, we performed ChIP-seq of ΔCTT.R3, a variant deficient in CBP/p300 binding and Aire condensate formation (Figs. 2d and 3d). Compared to WT Aire, ΔCTT.R3 strikingly exhibited a more dispersed genomic occupancy pattern, characterized by approximately tenfold more Aire-bound peaks (1,363 and 12,893 for WT Aire and ΔCTT.R3 ChIP peaks, respectively; Extended Data Fig. 7a) and relatively uniform ChIP-seq signal intensities compared to WT Aire (Fig. 4a). Moreover, the ChIP-seq signals of ΔCTT.R3 displayed weaker correlation to CBP/p300 ChIP-seq signals than did WT Aire (Fig. 4b, middle panels). These distinctions became more apparent when examining the top 500 WT-preferred versus ΔCTT.R3-preferred sites (as defined by the WT-to-ΔCTT.R3 ChIP-seq signal ratio in Fig. 4a, Supplementary Table 5); WT-preferred sites exhibited a pre-existing high density of CBP/p300 occupancy, whereas ΔCTT.R3-preferred sites were largely devoid of CBP/p300 (Fig. 4c-e). Analysis of Aire occupancy at SEs also showed a higher enrichment of WT Aire compared to ΔCTT.R3 (Extended Data Fig. 7b), indicating that CTT contributes to SE preference. Furthermore, CBP/p300 degrader dCBP-1 impaired WT Aire binding to SEs (Fig. 4f,g). A similar effect was also seen with the catalytic inhibitor A-485 (Fig. 4f,g), suggesting that Aire may preferentially home in on the catalytically active form of CBP/p300 that often form clusters and accumulate at active genetic loci^2,72,73^.

Altogether, these results demonstrate that Aire’s localization to H3K27ac-rich regions is mediated by Aire CTT interaction with CBP/p300 at these regions. Moreover, our findings revealed that Aire condensate formation, transcriptional activity and Aire’s genomic targeting are all dependent on CTT and CBP/p300. This strong association suggests a close coupling between Aire’s genomic targeting and polymerization for functional condensate formation.

### Aire’s preference for enhancer sites is amplified by CARD polymerization

To further investigate the relationship between Aire polymerization and genomic targeting, we performed ChIP-seq of ΔCARD, a variant with an intact CTT but impaired condensate formation and transcriptional functions (Fig. 2b and Extended Data Fig. 7c). We reasoned that if Aire polymerization, together with CBP/p300 binding, is essential for proper targeting, we would expect ΔCARD to fail to preferentially accumulate on CBP/p300-rich loci. On the other hand, if polymerization is a consequence of genomic targeting, ΔCARD would show similar preference for CBP/p300-rich loci as observed with WT Aire.

Compared to WT and ΔCTT.R3, ΔCARD showed an intermediate genomic targeting behavior (Fig. 4a-d). Specifically, ΔCARD’s genomic occupancy correlated with p300 occupancy less than WT Aire, but more than ΔCTT.R3 (Fig. 4b, middle vs. right panels). While ΔCARD favored WT-preferred sites over ΔCTT.R3-preferred sites, this preference was less pronounced in comparison to WT Aire (Fig. 4c,d). Similarly, ΔCARD’s SE localization was in between that of WT and ΔCTT.R3 (Extended Data Fig. 7b). Since both ΔCARD and ΔCTT lack condensate formation, but only ΔCTT lacks the preference for CBP/p300-rich sites, we infer that Aire’s target site selection is primarily driven by CTT–CBP/p300 interaction. However, the stronger target bias exhibited by WT Aire compared to ΔCARD suggests that CARD-mediated polymerization amplifies the CTT-driven genomic preference for CBP/p300-rich sites.

One plausible model to explain these findings is that Aire recruitment to CBP/p300-rich loci initiates CARD polymerization, facilitating the recruitment of additional Aire molecules and thereby enhancing Aire’s presence at the correct target site. The high density of CBP/p300 at Aire condensates and lack thereof in the absence of Aire (Fig. 1h and Extended Data Fig. 3a) suggests a feedback amplification that extends to CBP/p300, allowing for the recruitment of additional CBP/p300 molecules to the site of Aire nucleation. Altogether, while Aire’s targeting to CBP/p300-rich loci does not obligatorily require CARD polymerization, polymerization amplifies the target preference.

### PHD1 suppresses CARD polymerization

Our results thus far show that Aire requires CTT for condensate formation and this is likely mediated by CTT’s ability to direct Aire to CBP/p300-rich enhancers. However, the inability of ΔCTT to form any condensates was seemingly at odds with the fact that ΔCTT contains an intact CARD. Aire CARD can spontaneously polymerize *in vitro* and form condensates in cells^11^, and yet, is somehow unable to polymerize in the context of ΔCTT. This apparent discrepancy led us to hypothesize that Aire CARD polymerization is subject to regulatory mechanisms, and CTT-mediated targeting alleviates the suppression, ensuring controlled polymerization.

To identify the potential domain responsible for CARD regulation, we deleted individual domains in the AireΔCTT background, expecting that deletion of the CARD-suppressive domain would restore condensate formation. Deleting PHD1, but no other domains, restored condensate formation of AireΔCTT (Fig. 5a and Extended Data Fig. 8a), suggesting that PHD1 is the domain responsible for suppressing Aire polymerization in the absence of CTT. In line with the role of PHD1 in suppressing Aire polymerization, deletion of PHD1 in full-length Aire significantly increased the number of Aire condensates per nucleus (Fig. 5b).

**Fig. 5.**
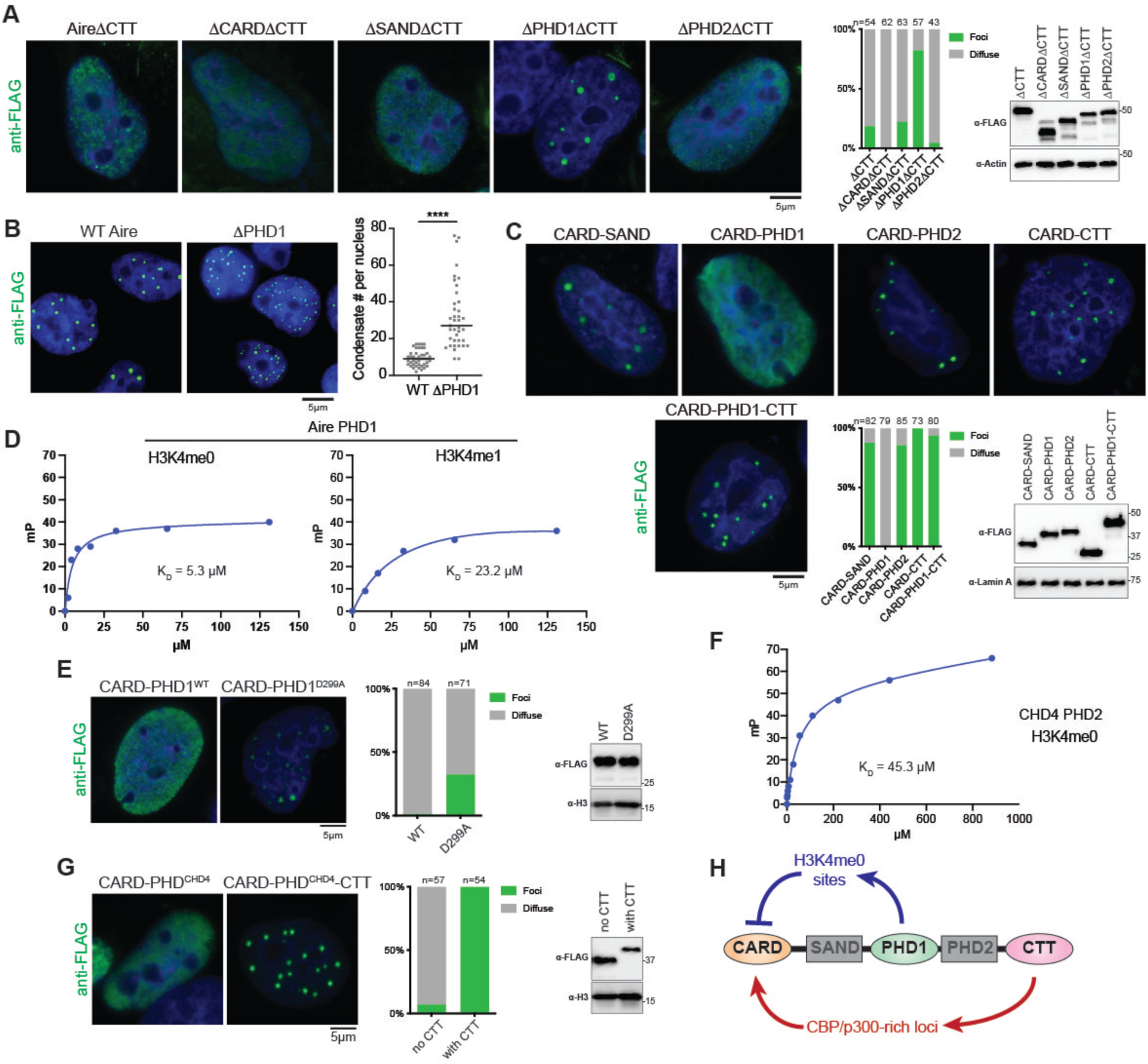
PHD1-mediated chromatin interaction suppresses CARD polymerization. (A) Representative immunofluorescence images of Aire domain deletion mutants in 4D6 cells. Cells were transfected with mouse Aire-FLAG expression plasmids 24 hrs prior to fixation. Middle: percentage of nuclei with Aire condensates vs. diffuse Aire staining. n represents the number of nuclei examined. Right: WB showing the levels of FLAG-tagged Aire variants. (B) Representative immunofluorescence images of WT Aire and ΔPHD1 in Dox-inducible stable 4D6 cells. Right: quantification of number of Aire condensates per nuclei. *p*-value (Mann-Whitney test) was calculated in comparison to WT Aire. \*\*\*\**p* < 0.0001. Experiments were done as in Fig. 2B. (C) Representative immunofluorescence images of mouse Aire CARD-fusion variants. Bottom middle: percentage of nuclei with Aire condensates vs. diffuse Aire staining. n represents the number of nuclei examined. Bottom right: WB showing the levels of FLAG-tagged Aire CARD-fusion variants. Experiments and analyses were done as in (A). (D) Binding of the indicated fluorescein-labeled histone H3 tail-derived peptides to increasing concentrations of mouse Aire PHD1. Binding is detected by a change in the fluorescence polarization (mP) of fluorescein. A representative experiment is shown for each peptide. The curves indicate the fit to a simple binding isotherm for the data sets shown. Data are representative of three independent experiments and presented as mean ± SD. H3K4me0 *K*_D_ = 5.25 ± 2.59 μM; H3K4me1 *K*_D_ = 23.18 ± 11.8 μM. (E) Representative immunofluorescence images of mouse Aire CARD-fusion variants. Experiments and analyses were done as in (A). (F) Binding of the fluorescein-labeled histone H3 tail-derived peptide (H3K4me0) to increasing concentrations of the CHD4 PHD2. Data are representative of three independent experiments and presented as mean ± SD. *K*_D_ = 45.3 ± 10.2 μM. (G) Representative immunofluorescence images of FLAG-tagged mouse Aire CARD fused with CHD4 PHD2 with and without Aire CTT. Experiments and analyses were done as in (A). (H) Schematic of the tightly coordinated interplay between Aire’s CARD, the histone-binding domain PHD1 and the activation domain CTT. As a negative regulator, PHD1 disperses Aire to numerous H3K4me0 sites across the entire genome, thereby “diluting” Aire and preventing CARD polymerization at inappropriate locations. At Aire-target sites, CTT recognizes CBP/p300-rich loci, leading to the local concentration of Aire and promotion of Aire CARD polymerization. All data are representative of at least three independent experiments.

To examine whether PHD1 is sufficient to suppress Aire CARD polymerization, we directly fused PHD1 with CARD and compared CARD-PHD1 with CARD fused with other domains of Aire. Only CARD-PHD1 showed diffuse staining, whereas other fusion constructs showed efficient condensate formation (Fig. 5c and Extended Data Fig. 8b). Furthermore, the PHD1-mediated suppression in the CARD-PHD1 construct was relieved by additional fusion of CTT (Fig. 5c), recapitulating the suppressive effect of PHD1 and stimulatory effect of CTT in full-length Aire condensate formation. These results suggest that PHD1 is the suppressor domain for CARD polymerization and the requirement for CTT in Aire condensate formation stems solely from PHD1-mediated suppression.

We next asked how PHD1 suppresses CARD polymerization. Because PHD1 is a histone binding domain with a specific preference for H3K4me0^33,34^ (*K*_D_ = 5.25 ± 2.59μM, Fig. 5d), we investigated whether PHD1 binding to H3K4me0 is important for inhibiting CARD polymerization. The D299A mutation in Aire PHD1, known to reduce the affinity for H3K4me0 and transcriptional activity of Aire^33,34,40^, compromised PHD1’s ability to suppress Aire CARD polymerization (Fig. 5e and Extended Data Fig. 8c). Additionally, fusing Aire CARD with another H3K4me0-specific PHD from an unrelated protein, CHD4^74^ (Fig. 5f), suppressed Aire CARD polymerization, while CTT restored it (Fig. 5g and Extended Data Fig. 8d). This suggests that Aire PHD1 suppresses CARD polymerization indirectly through H3K4me0 binding, rather than directly through a PHD1–CARD interaction. In keeping with this notion, nuclear condensate formation of a homologous CARD from another TR, Sp110, was also suppressed by fusing with Aire PHD1, but not with Aire PHD2 (Extended Data Fig. 8e). Considering that PHD1’s specificity for H3K4me0 is both necessary and sufficient for CARD suppression and given the widespread distribution of H3K4me0 throughout the genome, we speculate that PHD1 suppresses Aire polymerization by dispersing Aire across numerous genomic sites bearing H3K4me0. This dispersion would effectively dilute Aire and prevent its spontaneous nucleation. CTT, on the other hand, may counter this dilution effect by concentrating Aire at CBP/p300-rich loci, allowing target-specific polymerization (Fig. 5h).

### PHD1 suppression is necessary for transcriptionally active condensate formation

The role of PHD1 in suppressing CARD polymerization raised the question of whether PHD1 also negatively regulates Aire’s transcriptional function. Since condensate formation is necessary, more frequent condensate formation by ΔPHD1 could amplify or augment Aire’s function.

Alternatively, given that condensate formation is not sufficient for transcriptional function, it is also possible that ΔPHD1 condensates are transcriptionally inactive. RT-qPCR of Aire target genes showed that ΔPHD1 was transcriptionally inactive (Fig. 6a). Nascent RNA-FISH also showed significantly reduced transcriptional activity of ΔPHD1 despite forming more condensates per nucleus (Fig. 6b). Furthermore, ΔPHD1 condensates showed lower density of co-localized MED1 and CBP than WT Aire condensates (Fig. 6c). Similarly, other variants that bypassed PHD1-mediated regulation to form condensates (e.g. ΔPHD1ΔCTT, CARD-CTT) were transcriptionally inactive (Extended Data Fig. 9a,b). These results show that Aire condensates formed in the absence of PHD1-mediated suppression are transcriptionally inactive.

**Fig. 6.**
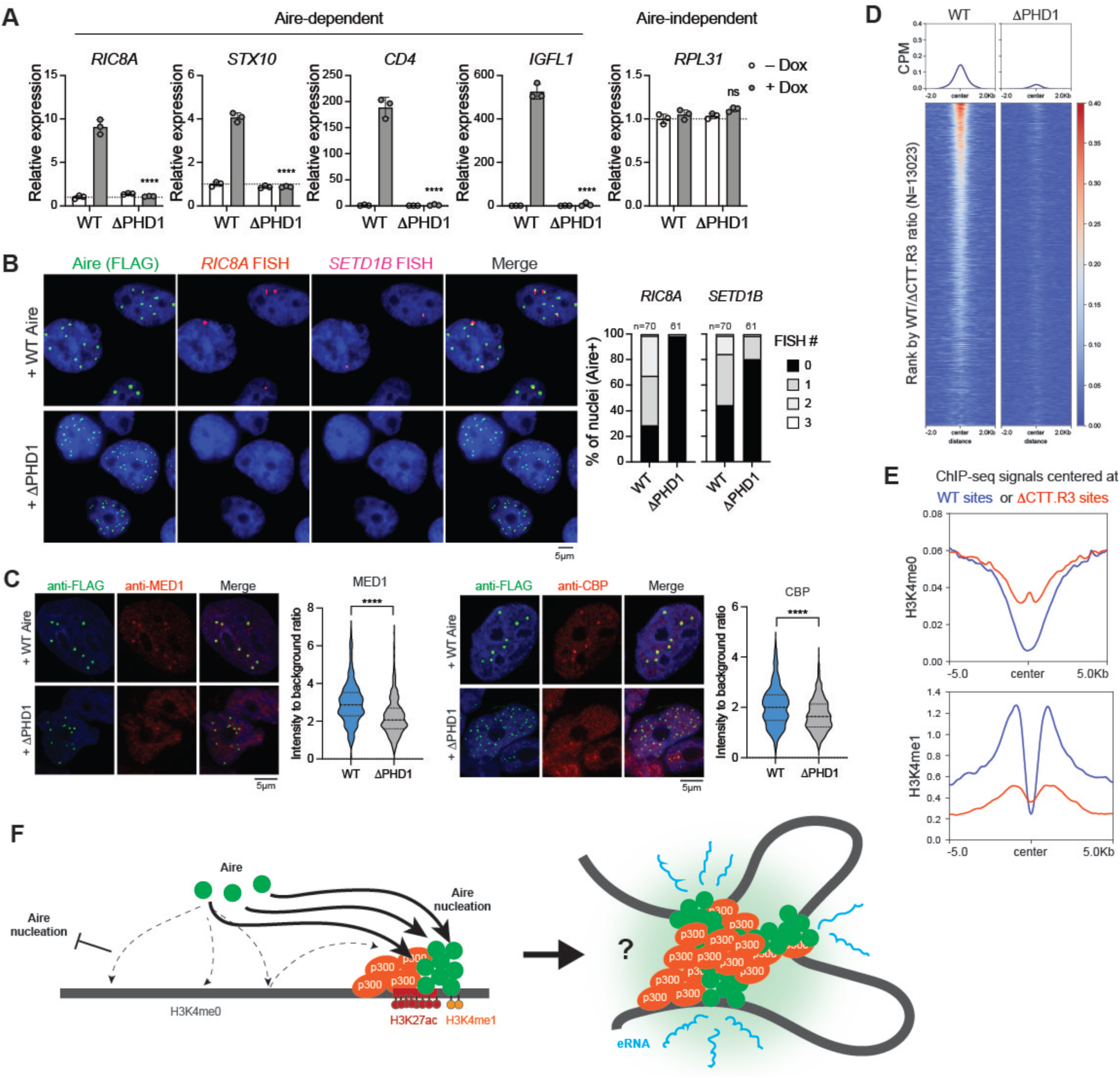
PHD1 cooperates with CTT to enable specific genomic targeting and formation of transcriptionally active condensates. (A) Transcriptional activity of WT Aire or ΔPHD1 mutant, as measured by the relative mRNA levels of Aire-dependent genes, *RIC8A*, *STX10*, CD4, and *IGFL1*, expressed in Dox-inducible stable 4D6 cells. An Aire-independent gene, *RPL31*, was also examined as a negative control. All genes were normalized against the internal control *RPL18*. Data are representative of at least three independent experiments and presented as mean ± SD, n = 3. *p*-values (two-tailed unpaired t-test) were calculated in comparison to WT Aire (+Dox). \*\*\*\**p* < 0.0001; *p* > 0.05 is not significant (ns). (B) Representative images showing IF of Aire-FLAG and nascent RNA-FISH of Aire-induced genes, *RIC8A* and *SETD1B*, after 24 hrs Dox-induction of WT Aire or ΔPHD1. Right: percent of nuclei that have 0, 1, 2 or 3 RNA-FISH foci (left: *RIC8A*; right: *SETD1B*). (C) Representative IF images of endogenous MED1 and CBP in 4D6 cells expressing Aire-FLAG WT or ΔPHD1 under the Dox-inducible promoter. Right: quantitation of the average intensities of MED1/CBP staining at Aire condensates. MED1/CBP intensities at Aire condensates were normalized to the average MED1/CBP intensity within the entire nucleus. ****p<0.0001 using a Mann-Whitney test. (D) Heatmaps of normalized ChIP-seq signal (CPM) for indicated proteins in 4D6 cells. Heatmaps are centered on all Aire peaks (n = 13,023) and ranked by the ratio of WT Aire over ΔCTT.R3 ChIP-seq signals as in Fig. 4A. Aire ChIP-seq signals shown were subtracted of background noise from corresponding input controls. For clarity, WT Aire ChIP-seq heatmap was reproduced from Fig. 4A. (E) Average H3K4me0 and H3K4me1 ChIP-seq profiles (normalized CPM) centered at WT-preferred sites versus DCTT.R3-preferred sites (top and bottom 500 peaks from Fig. 4A, respectively). Both H3K4me0 and H3K4me1 ChIP-seq experiments were performed on 4D6 cells prior to Aire expression. (F) Proposed mechanism for controlled Aire condensate assembly. Firstly, Aire employs PHD1’s affinity for ubiquitous unmethylated H3K4 to interact with chromatin in a distributive manner, preventing inappropriate CARD polymerization outside target enhancers. Secondly, Aire utilizes the CTT for specific targeting to CBP/p300-rich loci, countering PHD1’s dispersion effect. Targeted accumulation of Aire at CBP/p300-rich loci initiates a positive feedback loop, nucleating Aire CARD polymerization and recruiting additional Aire molecules, depleting dispersed Aire throughout the genome. Aire polymerization also recruits additional CBP/p300 molecules and connects multiple Aire-bound loci, ultimately leading to the formation of condensates highly enriched with Aire and CBP/p300. This conglomerate of Aire and CBP/p300 creates a potent environment capable of activating eRNA transcription without necessarily altering the levels of H3K27ac or chromatin accessibility. In summary, the coordinated action of PHD1-mediated “regulation-by-dispersion” and CTT-mediated targeting is essential for functional Aire condensate formation.

To understand why ΔPHD1 is transcriptionally inactive, we analyzed ΔPHD1’s interaction with chromatin. ΔPHD1 ChIP-seq revealed only 7 ΔPHD1-occupied peaks, as opposed to 1,363 peaks with WT Aire, all with the same peak calling and analysis pipeline (Extended Data Fig. 7a). When comparing ChIP signals of ΔPHD1 vs. WT Aire, ΔPHD1 showed remarkably lower signal at nearly all Aire-bound sites (Fig. 6d), indicating that PHD1 deletion completely abolished Aire’s chromatin binding.

To confirm that ΔPHD1 is impaired in chromatin interaction, we performed a nuclear fractionation analysis, where solubility of chromatin and chromatin-associated proteins were monitored in the presence and absence of MNase. If a protein binds chromatin, it would be present in the insoluble pellet (P) together with histone, until MNase releases nucleosomes and associated factors into the soluble fraction (S). Both WT and ΔPHD1 co-fractionated with histones in the P fraction in the absence of MNase treatment (Extended Data Fig. 9c). Upon MNase treatment, a subset of WT Aire was released to the S fraction, indicating WT Aire associates with chromatin. By contrast, ΔPHD1 existed solely in the P fraction independent of MNase treatment, suggesting an impairment in chromatin binding (Extended Data Fig. 9c), indicating the formation of insoluble aggregates that are detached from chromatin. Thus, the defect in transcriptional activity of ΔPHD1 condensate is likely due to the lack of chromatin tethering. These results show that while CTT and CARD can modulate Aire’s target specificity, PHD1 is absolutely necessary for Aire’s association with chromatin.

How does PHD1, with its preference for H3K4me0, promote Aire’s interaction with chromatin at CBP/p300-rich enhancers where H3K4 is usually methylated^43,44^? One possibility is that the role of PHD1 in facilitating Aire’s chromatin localization may be an indirect consequence of suppressing aberrant polymerization of Aire. Once spontaneously polymerized, Aire may be in a conformation that is no longer competent for chromatin binding. Alternatively, PHD1 may directly participate in target recognition by binding to lower affinity substrates, such as H3K4me1 (Fig. 5d), that are nearby CBP/p300 clusters. Indeed, histone ChIP-seq showed that Aire’s target sites (WT-preferred) displayed high levels of H3K4me1 nearby, while being depleted of H3K4me0 (Fig. 6e, also see Extended Data Fig. 9d for other histone marks). In contrast, Aire’s non-target sites, such as ΔCTT.R3-preferred sites, had relatively higher levels of H3K4me0, but lower levels of H3K4me1 (Fig. 6e). Regardless of the specific mechanism, our results unambiguously demonstrate that PHD1-mediated suppression of spontaneous CARD polymerization serves as an important intermediate step for the eventual formation of transcriptionally active Aire condensates.

## Discussion

Recent studies have highlighted the importance of biomolecular condensates in transcriptional regulation^1–4^. While studies on transcriptional condensates have largely focused on identifying domains for multimerization, how multimerization is regulated to ensure proper localization and prevent aberrant condensate formation is relatively poorly understood. Such regulation is particularly important for CARD-containing TRs, which can spontaneously polymerize. The importance of controlled polymerization extends beyond TRs; numerous CARD-containing proteins involved in cytoplasmic cell death and inflammatory signaling pathways also rely on stringent regulatory mechanisms, although our understanding of these mechanisms also remains limited^17,18^.

We here reveal a multi-layered regulatory mechanism that enables Aire to form condensates at appropriate target sites, namely CBP/p300-rich enhancers. First, Aire utilizes PHD1 to limit spontaneous polymerization, and this suppression is mediated through its ability to bind H3K4me0. Loss of PHD1 results in more frequent and uncontrolled formation of Aire condensates that are detached from chromatin and transcriptionally inactive. We propose that by interacting with H3K4me0 ubiquitously present throughout the genome, PHD1 allows Aire to bind chromatin in a distributive manner, preventing CARD polymerization. Second, CTT counters this PHD1-mediated suppression by guiding Aire to the CBP/p300-rich loci and nucleating CARD polymerization.

The precise coupling mechanism between Aire’s genomic targeting and CARD polymerization remains unclear, but one possibility is that Aire’s focused recruitment to CBP/p300 clusters increases the local concentration of Aire, overriding PHD1-mediated “dilution” effect and facilitating CARD nucleation. Once CARD polymerizes, an Aire nucleation site then acts as a sink to recruit more Aire and CBP/p300 molecules, setting up a positive feedback loop for transcriptional condensate assembly and local transcriptional activation (Fig. 6f). Importantly, CTT’s role in stimulating Aire polymerization is contingent upon PHD1-mediated suppression, suggesting that CTT does not directly participate in polymerization as with other ADs^2,5,6^. In essence, PHD1-mediated chromatin anchoring makes Aire polymerization strictly dependent on CTT and restricted to activated CBP/p300-rich environments, such as SEs.

Once Aire forms transcriptional condensates, how does Aire activate its genomic binding sites and how do these molecular activities relate to Aire’s ultimate function of regulating PTA expression? We found that Aire condensates, although nucleated at CBP/p300-rich loci, became more enriched with CBP/p300 (as evidenced by increased CBP/p300 at Aire condensates) and activated eRNA transcription, yet with minimal increase in H3K27ac or chromatin accessibility. These results suggest that Aire may harness CBP/p300’s scaffolding functions^75,76^, recruiting additional coactivators, bringing together multiple Aire-bound loci (Fig. 6f), and altering cohesin-mediated chromatin architecture^77^. From this perspective, the role of Aire may be to augment pre-existing enhancers by consolidating multiple distinct enhancers and cementing an enhancer landscape (Fig. 6f). These functions align with recent findings that one of Aire’s primary roles is to assist and amplify the activities of lineage-defining TFs that are responsible for mediating PTA expression^29–31,52^. In the non-native cellular environment of mTECs, these lineage-defining TFs may need Aire’s assistance in shaping the chromatin landscape. Once such landscape is optimized for PTA expression, Aire may become dispensable, explaining the loss of Aire expression in later stages of mTEC development^29–31,52^. In summary, our mechanistic insights provide a foundation for investigating how Aire activates the Aire-bound loci and regulates PTA expression.

---

**Extended Data Fig. 1.**
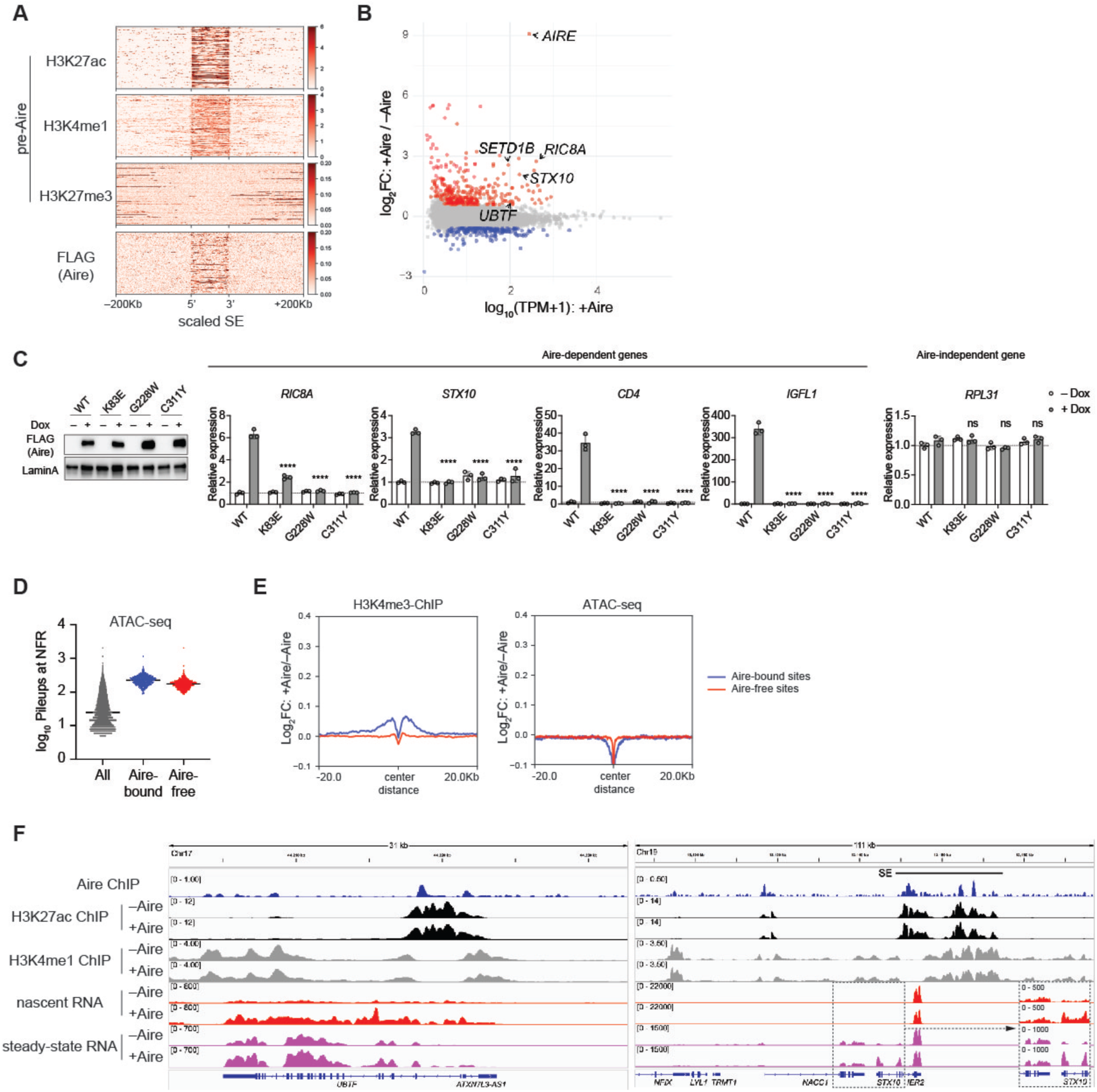
Thymic epithelial cell line (4D6) with Dox-inducible Aire recapitulates Aire transcriptional activity and functions. (A) Aire ChIP heatmaps of normalized ChIP-seq signals (Counts Per Million, CPM) for indicated proteins 200Kb up- or downstream of H3K27ac-delimited super-enhancers (n = 529). Histone marks are shown in 4D6 cells without Aire expression (pre-Aire). (B) Gene expression (transcripts per million, TPM) in Aire-expressing 4D6 cells (+Aire) versus expression changes between cells with and without Aire (+Aire/–Aire). Steady-state bulk RNA-seq was performed on Dox-inducible Aire-expressing 4D6 cells without or with 24 hr Dox treatment. Red, genes upregulated by Aire (>1.5-fold, FDR<0.05); blue, genes downregulated by Aire (<1.5-fold, FDR<0.05). FC, fold-change. FDR, false discovery rate. (C) Transcriptional activity of WT Aire or APS-1 mutants, as measured by the relative mRNA levels of Aire-dependent genes, *RIC8A*, *STX10*, *CD4* and *IGFL1*, in Dox-inducible 4D6 cells. An Aire-independent gene, *RPL31*, was also examined as a negative control. All genes were normalized against the internal control *RPL18*. Data are presented as mean ± SD, n = 3. *p*-values (one-way ANOVA with Dunnett’s multiple comparisons test) were calculated in comparison to WT Aire (+ Dox). \*\*\*\**p* < 0.0001; *p* > 0.05 is not significant (ns). Left: a western blot (WB) showing the nuclear levels of FLAG-tagged Aire proteins compared to endogenous levels of Lamin A. (D) Pileup counts at all nucleosome-free regions (NFRs), Aire-bound or Aire-free NFRs that showed similar ATAC-seq signals. n = 215039, 542 and 658, respectively. (E) Aire-induced changes in H3K4me3 ChIP or ATAC-seq 20 kb upstream or downstream of Aire-bound and Aire-free NFRs, n = 542 and 658, respectively. (F) Genome browser views of normalized ChIP-seq and RNA-seq profiles at exemplar Aire-bound sites in 4D6 cells. Numbers to the left indicate the ranges of normalized reads for RNA-seq or CPM for ChIP-seq. SE, super-enhancer.

**Extended Data Fig. 2.**
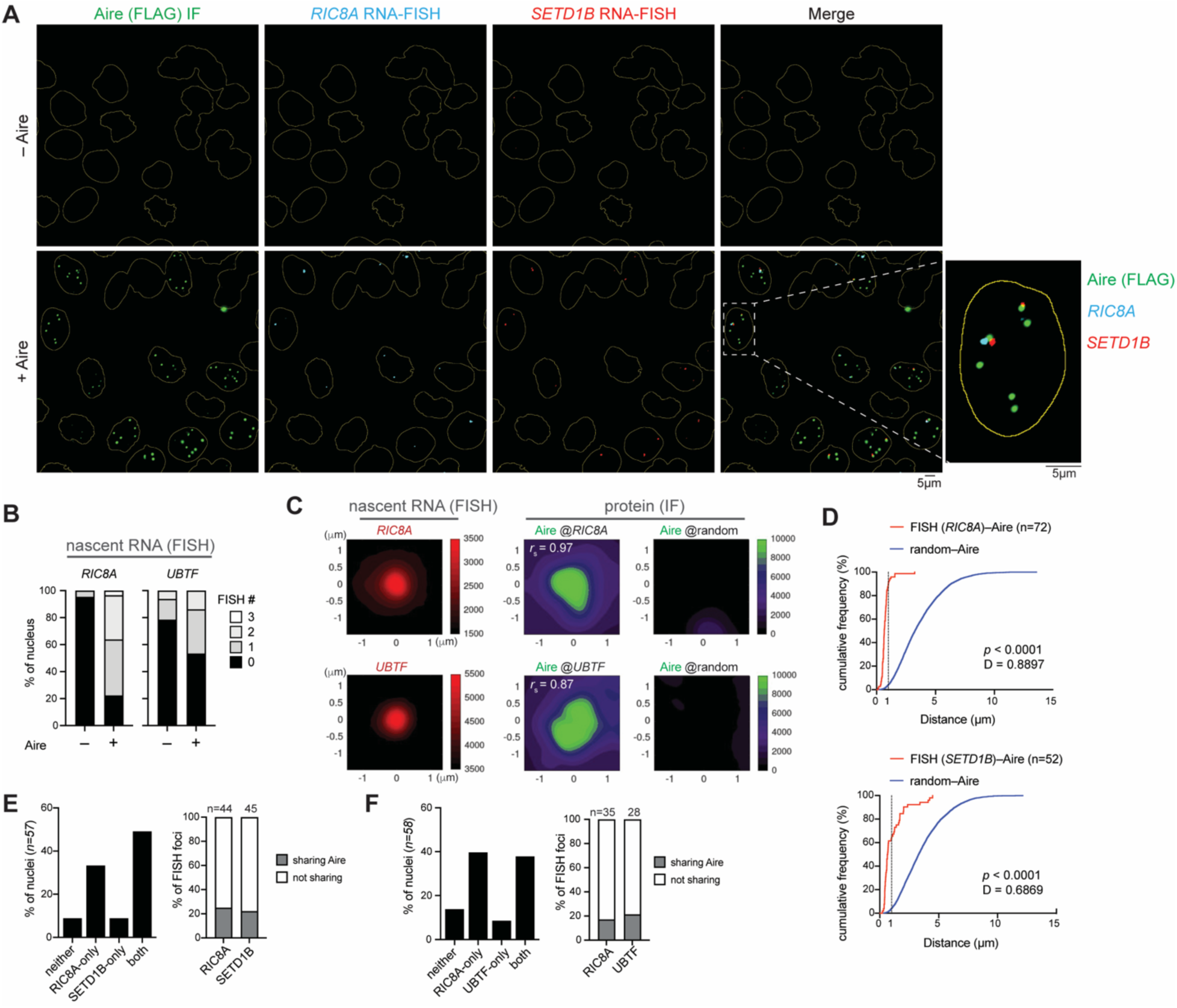
Aire links inter-chromosomal genomic loci into transcriptionally active condensates. (A) Nascent RNA-fluorescence in situ hybridization (FISH) coupled with immunofluorescence (IF) images of 4D6 cells expressing Aire-FLAG under a Dox-inducible promoter. Cells were not (–Aire) or were induced with 1 µg/ml Dox (+Aire) for 24 hrs before staining with FISH probes and anti-FLAG. RNA-FISH probes were designed to hybridize to the intronic regions of Aire-dependent targets. Yellow outlines represent the boundaries of nuclei from DAPI staining. Bottom right: zoomed-in view of the white-dashed box. For clarity, the zoomed-in image of a nuclei with *RIC8A* and *SETD1B* RNA-FISH foci sharing the same Aire focus was reproduced from Fig. 1B. (B) Quantitation of percent of nuclei that have 0, 1, 2 or 3 RNA-FISH foci (left: *RIC8A*; right: *UBTF*). A total of 65 and 58 nuclei were examined in cells treated without (–Aire) and with Dox (+Aire), respectively. (C) Quantitation of average intensity signals from RNA-FISH combined with IF of Aire-FLAG in 4D6 cells. Shown are average signals of RNA-FISH (left), Aire IF centered on FISH foci (center) and Aire IF centered on randomly selected nuclear positions (right). *r*_s_ denotes the Spearman’s correlation coefficient between RNA-FISH or IF signals. (D) Cumulative distribution of the minimum center-to-center distances between Aire condensates and observed RNA-FISH foci (FISH–Aire) or between Aire condensates and simulated random positions (random–Aire). D indicates Kolmogorov-Smirnov test D statistic. *p*, *p*-value. While RNA-FISH and Aire condensates often did not show perfect concentric colocalization, this is in line with recent studies showing that active genetic elements can be partially (within ∼1 µm), rather than completely, overlapping with the target genes by imaging^3–5^. The close association between Aire condensates and the target gene RNA-FISH foci therefore is consistent with the notion that Aire condensates directly activate transcription. (E) Quantitation of percent of nuclei harboring RNA-FISH foci. Left: statistics of *RIC8A* and *SETD1B* RNA-FISH foci from (A) and Fig. 1B. Right: frequency of RNA-FISH foci sharing Aire condensates, when measuring within nuclei showing RNA-FISH foci of both genes. (F) Quantitation of percent of nuclei harboring RNA-FISH foci. Left: statistics of *RIC8A* and *UBTF* RNA-FISH foci from (B). Right: frequency of RNA-FISH foci sharing Aire condensates, when measuring within nuclei showing RNA-FISH foci of both genes.

**Extended Data Fig. 3.**
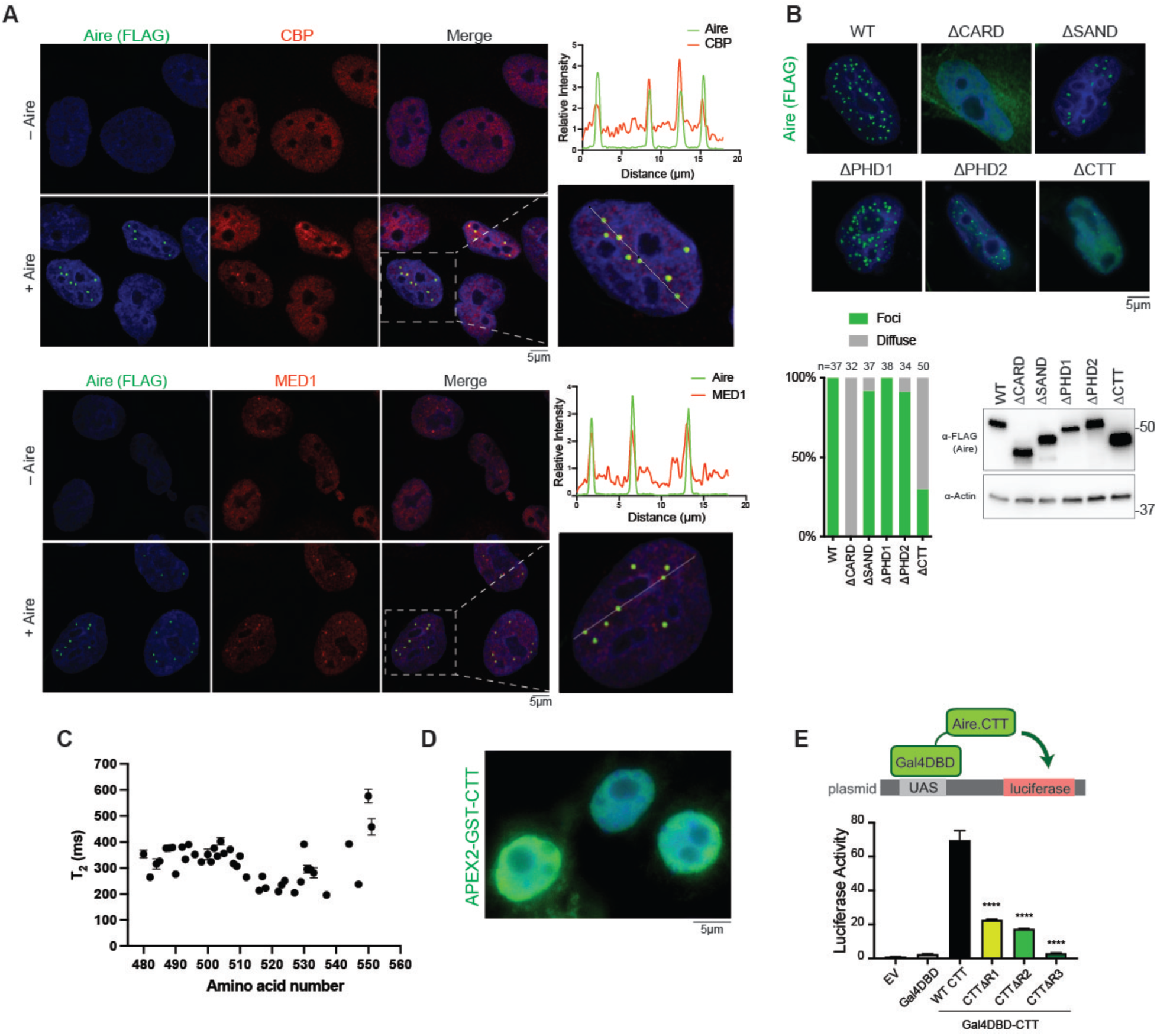
Aire condensates represent sites of active transcription and require Aire CTT. (A) Representative immunofluorescence images of endogenous CBP and MED1 in 4D6 cells before and after Aire-FLAG expression. Cells were not (– Aire) or were induced with 1 µg/ml Dox (+ Aire) for 24 hrs before immunostaining with anti-FLAG, anti-CBP and anti-MED1. Right: zoomed-in views of nuclei enclosed within white-dashed boxes along with measured fluorescence intensities across drawn solid white lines. (B) Representative fluorescence images of Aire WT and domain deletion variants in 4D6 cells. Cells were transfected with mouse Aire-FLAG expression plasmids 24 hrs prior to fixation. Bottom left: percentage of nuclei with Aire condensates versus diffuse Aire staining. n represents the number of nuclei examined. Bottom right: WB showing the levels of the indicated proteins. (C) 15N relaxation time (T_2_) of Aire CTT (340 µM) recorded at 700 MHz and 15°C. The averaged T_2_ is 0.319 s (corresponding relaxation rate (R_2_) of 3.13 s-1). (D) Representative fluorescence image of APEX2-GST-CTT in 293T. Cells were transfected with a plasmid constitutively expressing mouse Aire CTT fused with APEX2-GST for 24 hrs prior to fixation and staining with anti-FLAG. (E) TAD-like activity of mouse Aire CTT and various CTT deletion mutants. Top: schematic of the assay. CTT was fused with Gal4 DNA-binding domain (Gal4DBD), which binds upstream activation sequences (UAS) and controls the expression level of the reporter luciferase. The effect of Gal4DBD-CTT on the reporter activity was measured in 4D6 cells. Bottom: luciferase activities were shown relative to that of empty vector (EV)-transfected cells. Data are representative of at least three independent experiments and presented as mean ± SD, n = 3. *p*-values (one-way ANOVA with Dunnett’s multiple comparisons test) were calculated in comparison to WT CTT, \*\*\*\**p* < 0.0001.

**Extended Data Fig. 4.**
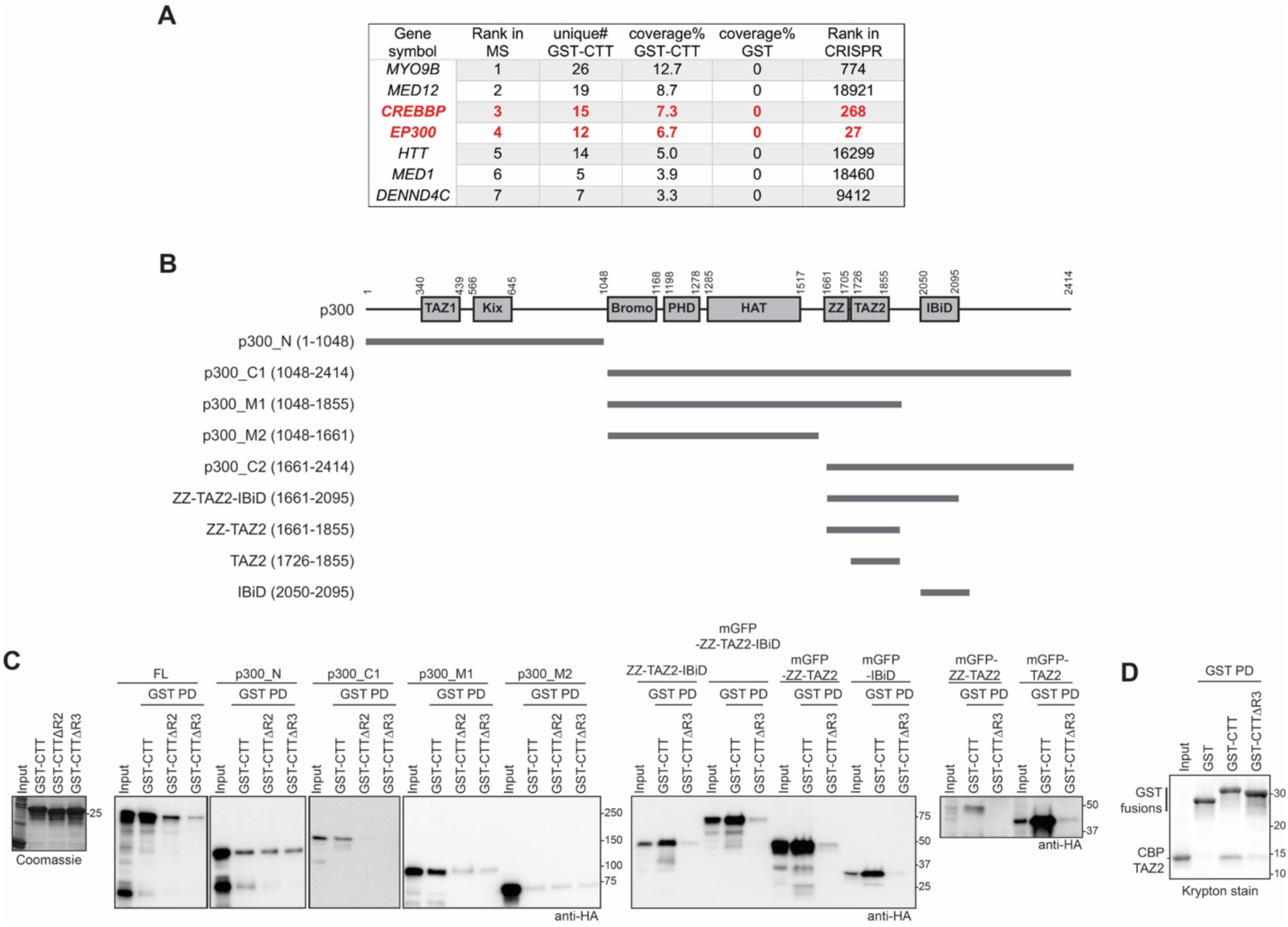
CBP/p300 directly interact with Aire CTT. (A) LC-MS/MS analysis results of the GST-CTT pull-down in Fig. 3C. A ∼250 kDa band in the GST-CTT-bound fraction and the equivalent region in the GST control were analyzed. Also see Supplementary Table 4 for more details. (B) Domain architecture of p300 and the truncation variants used in (C). Note that the homolog CBP has the same domain architecture. (C) His_6_-GST-CTT variant pull-downs of HA-tagged p300 transiently expressed in 293T nuclear extracts. Far left: SDS-PAGE gel of captured GST-CTT variant proteins captured onto glutathione beads used for the pull-down assay. Right: WBs of HA-p300 variants pulled-down with GST-CTT WT vs. GST-CTTΔR2 or GST-CTTΔR3. (D) His_6_-GST-CTT variant pull-downs of recombinantly expressed and purified CBP TAZ2.

**Extended Data Fig. 5.**
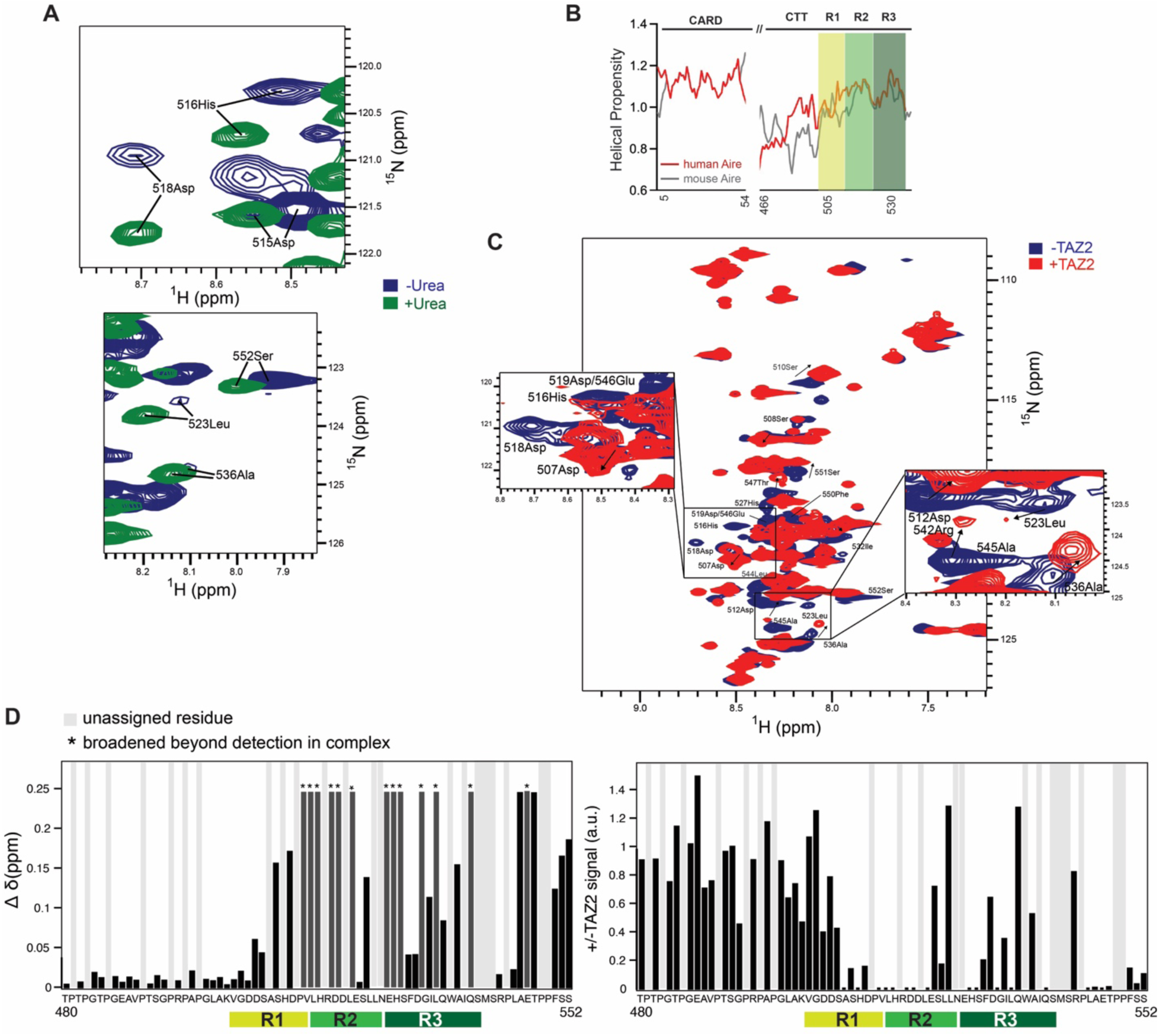
Aire CTT R2 and R3 bind CBP TAZ2. (A) NMR spectroscopy of mouse Aire CTT (aa 480-552) in the absence or presence of the chaotrope urea to identify regions with structural propensity, as measured by exchange broadening. 2D-^15^N-HSQC of ^15^N^13^C-labeled Aire CTT was recorded at 15°C (288K) on an 800 MHz spectrometer equipped with a cryoprobe. To assign the CTT regions with exchange broadening, urea was added to ^15^N^13^C-labeled Aire CTT. Aire residues in both panels are severely exchange broadened in the absence of urea (blue), suggesting secondary structure propensity. Upon addition of urea (green), the structural propensity is lost, and the highlighted peaks gain in intensity, allowing for their assignment. The results show that, in the absence of TAZ2, isolated CTT is largely disordered, as evident from low dispersion in the proton chemical shift and secondary chemical shifts. R2 and R3, on the other hand, alternate between disordered and alpha helical characteristics, as evidenced by strong peak broadening. The helical propensities of R2 and R3 are further supported by secondary structure predictions (B) and AlphaFold modeling (Fig. 3E). (B) Helical propensity of CTT from human and mouse Aire (ProtScale^6^, using Chou & Fasman parameters). Aire CARD, which has high α-helical propensity, was used in comparison. (C) NMR spectroscopy of ^15^N^13^C-labeled mouse Aire CTT (aa 480-552) in the absence or presence of CBP TAZ2. 2D-^15^N-HSQC of ^15^N^13^C-labeled Aire CTT alone (blue) and with 1:1.75 molar ratio of ^15^N^13^C-labeled Aire CTT:CBP TAZ2 (red), recorded at 15°C (288K) on an 800 MHz spectrometer equipped with a cryoprobe. Close-ups at a lower contour level show the extent of peak broadening and chemical shift perturbation upon complex formation in select regions. (D) Plots of detected chemical shift perturbations (CSPs) Δ*δ* [calculated as Δ*δ*=sqrt(0.14∗Δ*N*^2^+Δ*H*^2^)]^7^ and the signal loss upon addition of CBP TAZ2 (1:1.75 molar ratio of Aire CTT:CBP TAZ2). Mouse Aire CTT residues marked with an asterisk were broadened beyond detection in the Aire−CBP complex.

**Extended Data Fig. 6.**
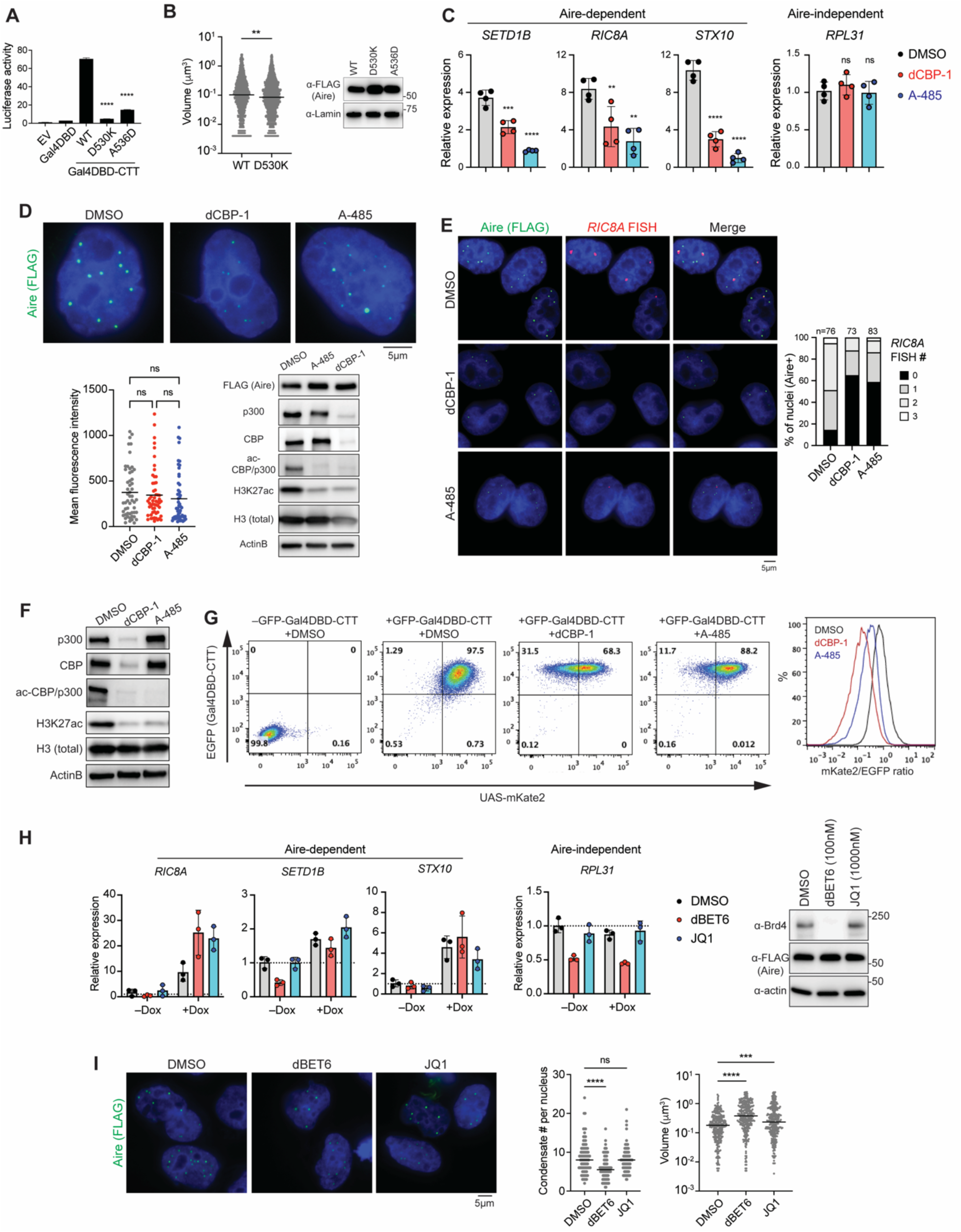
CBP/p300 are essential for Aire transcription condensate formation. (A) TAD-like activity of WT Aire CTT and point mutants as measured by the Gal4-luciferase assay in Extended Data Fig. 3D. Mouse Aire CTT was used. Data are presented as mean ± SD, n = 3. *p*-values (one-way ANOVA with Dunnett’s multiple comparisons test) were calculated in comparison to Gal4DBD-CTT WT (B) Quantification of WT vs. D530K Aire condensate volumes within all Aire-expressing nuclei examined in Fig. 3I. Right: WB showing the levels of WT Aire and mutants in transiently transfected cells. (C) Transcriptional activity of Aire in the presence of dCBP-1 (0.25 µM) or A-485 (3 µM) as measured by nascent RNA RT-qPCR of *SETD1B*, *RIC8A* and *STX10*. Cells were treated as in Fig. 3J. Data are presented as mean ± SD, n = 4. *p*-values (one-way ANOVA with Dunnett’s multiple comparisons test) were calculated in comparison to DMSO-treated cells (+ Dox for 24hr). \*\**p* < 0.01; \*\*\**p* < 0.001; \*\*\*\**p* < 0.0001; *p* > 0.05 is not significant (ns). (D) Representative immunofluorescence images of FLAG-tagged WT Aire in DMSO–, dCBP-1– or A-485–treated 4D6 cells. Bottom left: mean fluorescence intensity of nuclei examined. *p*-values (Kruskal-Wallis test with Dunn’s multiple comparisons test) were calculated. *p* > 0.05 is not significant (ns). Bottom right: WB showing the levels of CBP/p300 and their acetylation targets after drug treatment. Experiments were done as in Fig. 3K. (E) Representative images showing IF of Aire-FLAG and nascent RNA-FISH of Aire-induced gene *RIC8A* in the presence of dCBP-1 and A-485 in 4D6 cells. Right: percent of nuclei that have 0, 1, 2 or 3 RNA-FISH foci; n represents the number of nuclei examined for each sample. Drug treatments of cells were done as in Fig. 3K. (F) WB showing the levels of indicated proteins post-24hr treatment of the p300/CBP degrader dCBP-1 (0.25 µM) or the catalytic inhibitor A-485 (3 µM) in comparison to vehicle control DMSO in 4D6 cells. (G) Flow cytometry plots showing expression levels of the mKate2 reporter versus GFP-Gal4DBD-CTT (left) or the distribution of their ratios (right) in 4D6 cells. Cells were treated with Dox to induce GFP-Gal4DBD-CTT expression in the presence of DMSO, 0.25 µM dCBP-1 or 3 µM A-485 for 24 hrs prior to flow cytometry examination. (H) Transcriptional activity of Aire in the presence of dBET6 or JQ1, as measured by nascent RNA RT-qPCR of *SETD1B*, *RIC8A* and *STX10*. Dox, dBET6 (100 nM) or JQ1 (1000 nM), and 5’-EU were added to cell culture 24 hrs, 4hrs, and 0.5 hr prior to RNA extraction, respectively. Data are presented as mean ± SD, n = 3. Right: WB showing the levels of Brd4 and Aire-FLAG after drug treatment. (I) Representative immunofluorescence images of FLAG-tagged WT Aire in DMSO–, dBET6– or JQ1–treated 4D6 cells. Middle: quantification of number of Aire condensates per nuclei. Right: quantification of Aire condensate volumes within cells treated with different Brd4 inhibitors. *p*-values (Kruskal-Wallis test with Dunn’s multiple comparisons test) were calculated in comparison to DMSO-treated cells. \*\*\**p* < 0.001; \*\*\*\**p* < 0.0001; *p* > 0.05 is not significant (ns).

**Extended Data Fig. 7.**
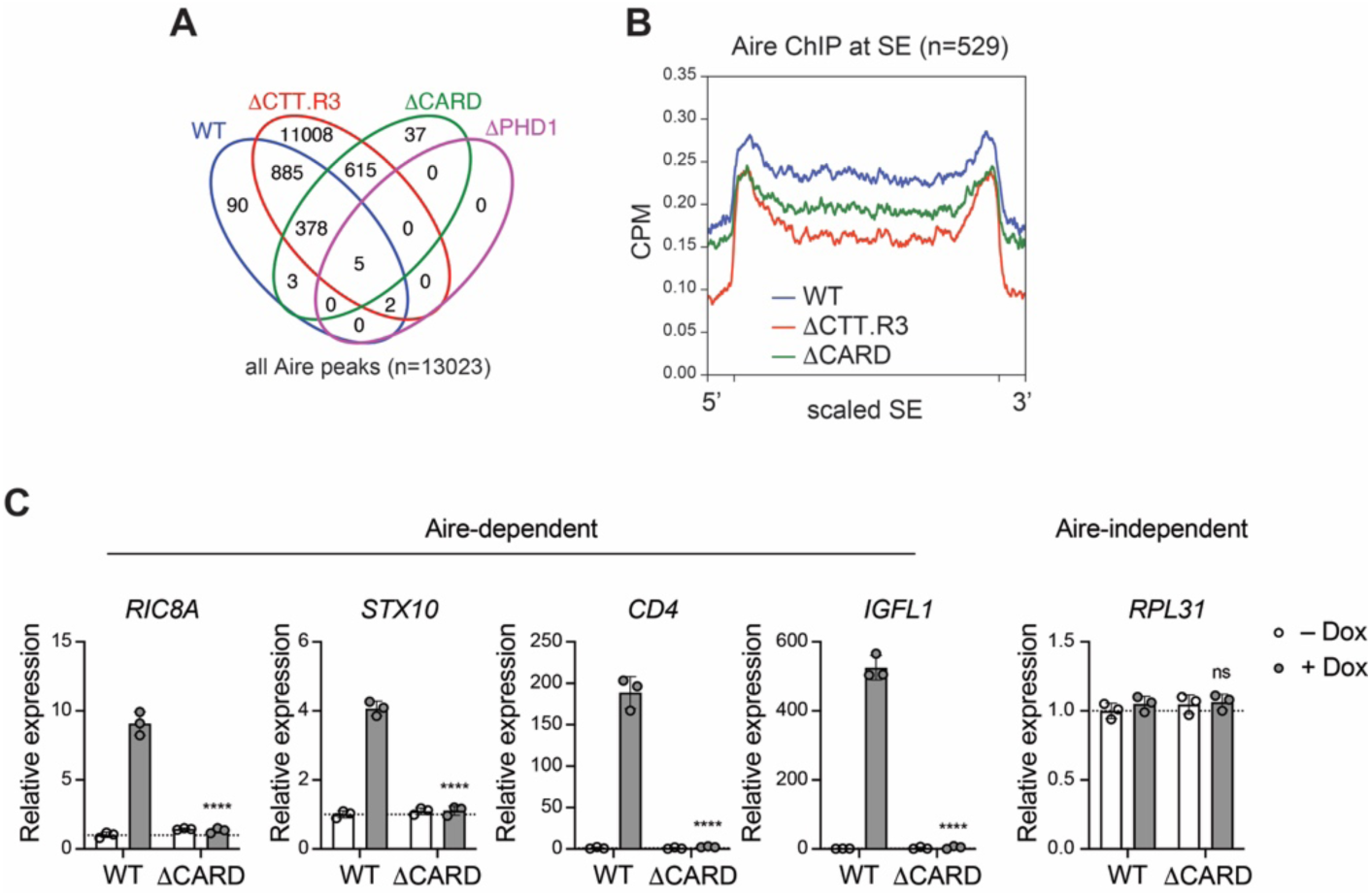
CARD is essential for robust Aire localization to super enhancers and mediating transcriptional activity. (A) Venn diagram showing all peaks identified from WT vs. mutant Aire ChIP-seq. (B) Average WT Aire, DCTT.R3, and DCARD ChIP-seq profiles (CPM) at H3K27ac-delimited super-enhancers (SEs, n = 529, defined in Fig S1A). *p*-values were calculated using Wilcoxon rank sum test (*p* < 2.2×10^-16^ for both WT vs. ΔCTT.R3 and WT vs. ΔCARD). (C) Transcriptional activity of WT Aire or ΔCARD mutant, as measured by the relative mRNA levels of Aire-dependent genes, *RIC8A*, *STX10*, CD4, and *IGFL1*, in Dox-inducible 4D6 cells. An Aire-independent gene, *RPL31*, was also examined as a negative control. Data are representative of at least three independent experiments and presented as mean ± SD, n = 3. *p*-values (two-tailed unpaired t-test) were calculated in comparison to WT Aire (+Dox). \*\*\*\**p* < 0.0001; *p* > 0.05 is not significant (ns).

**Extended Data Fig. 8.**
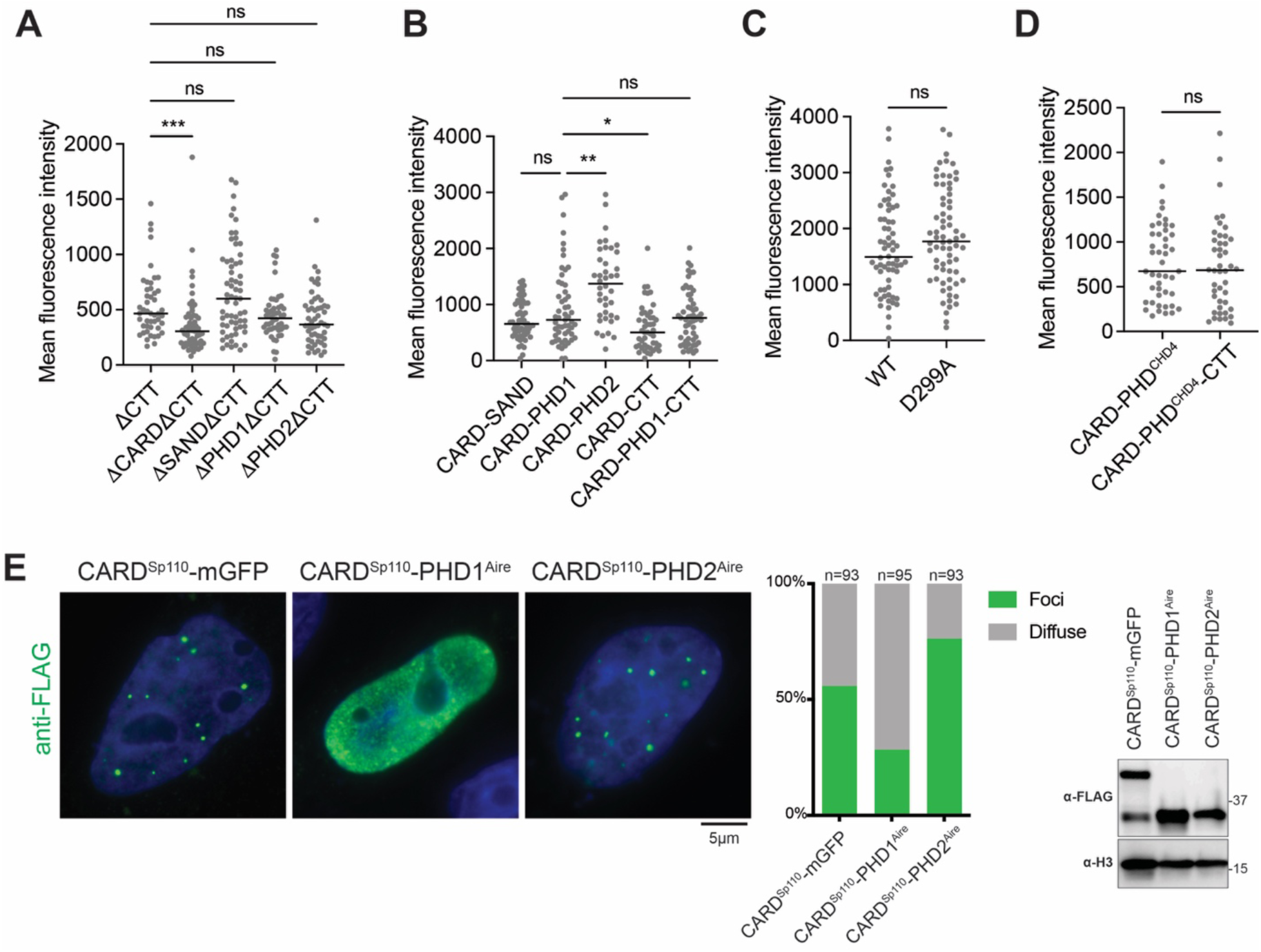
PHD1 binding to H3K4me0 is sufficient to inhibit Aire CARD polymerization. (A-D) Mean fluorescence intensity of nuclei examined corresponding to IF images in Fig. 5A, 5C, 5E, and 5G, respectively. *p*-values (Kruskal-Wallis test with Dunn’s multiple comparisons test for Fig.s A-B and Mann-Whitney test for Fig. C-D) were calculated in comparison to the control groups indicated in each graph. \**p* < 0.05; \*\**p* < 0.01; \*\*\**p* < 0.001; *p* > 0.05 is not significant (ns). (E) Representative immunofluorescence images of FLAG-tagged Sp110 CARD fused with mGFP (monomeric GFP), or PHD1 or PHD2 from mouse Aire. Experiments and analyses were done as in Fig. 5A.

**Extended Data Fig. 9.**
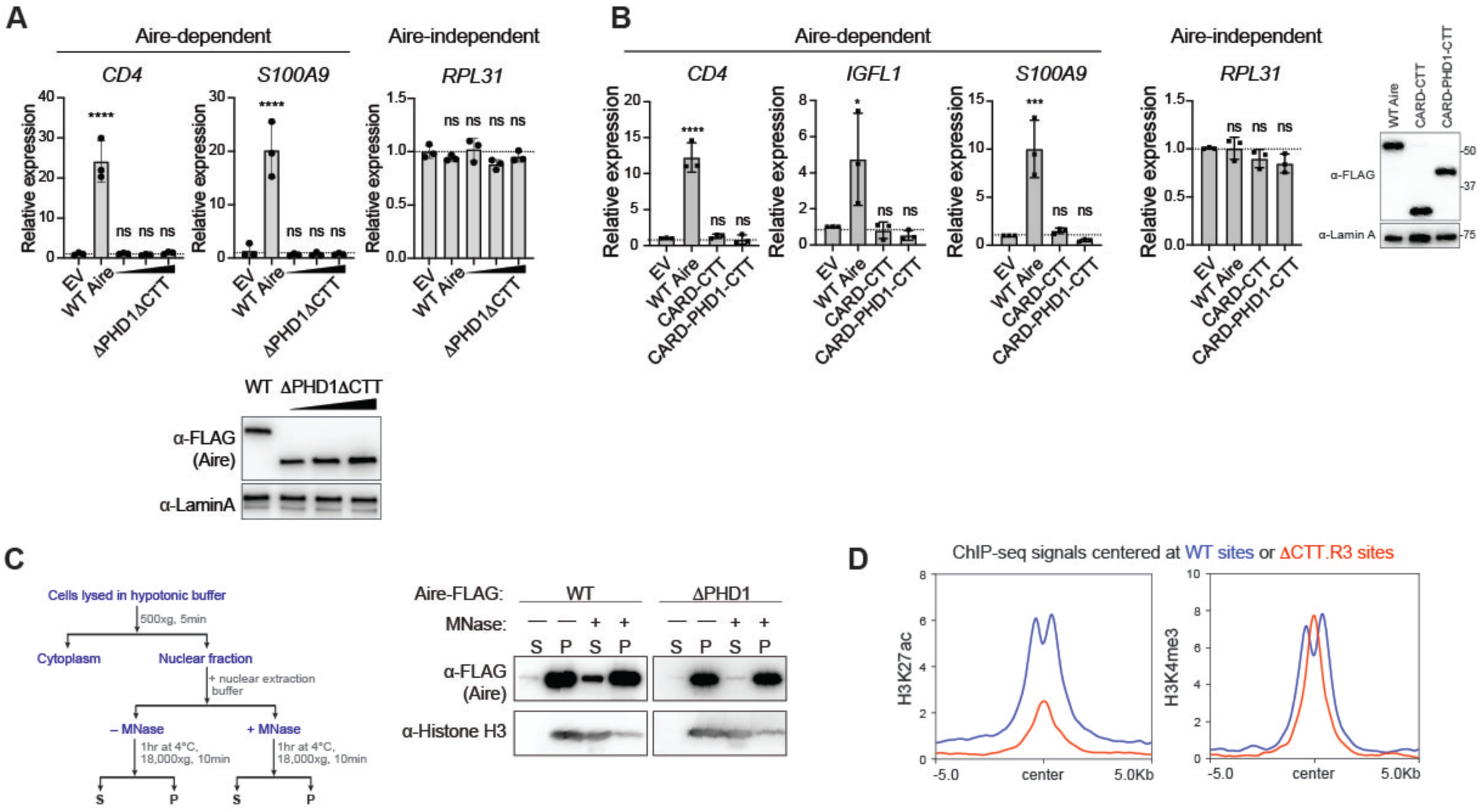
PHD1’s ability to bind H3K4me0 is required for the chromatin anchoring of Aire. (A) Transcriptional activity of WT Aire or ΔPHD1DCTT mutant, as measured by the relative mRNA levels of Aire-dependent genes, CD4, and *S100A9*, in 4D6 cells transiently transfected with plasmids expressing mouse Aire variants. An Aire-independent gene, *RPL31*, was also examined as a negative control. Data are representative of at least three independent experiments and presented as mean ± SD, n = 3. *p*-values (one-way ANOVA with Dunnett’s multiple comparisons test) were calculated in comparison to empty-vector transfected cells (EV). \*\*\*\**p* < 0.0001; *p* > 0.05 is not significant (ns). (B) Transcriptional activity of WT Aire or CARD-fusions, as measured by the relative mRNA levels of Aire-dependent genes, CD4*, IGFL1* and *S100A9*, in 4D6 cells transiently transfected with plasmids expressing mouse Aire variants. An Aire-independent gene, *RPL31*, was also examined as a negative control. Data are representative of at least three independent experiments and presented as mean ± SD, n = 3. *p*-values (one-way ANOVA with Dunnett’s multiple comparisons test) were calculated in comparison to empty-vector transfected cells (EV). \*\*\*\**p* < 0.0001; \*\*\**p* < 0.001; *p* > 0.05 is not significant (ns). (C) Chromatin fractionation analysis of Aire WT and ΔPHD1. Left: schematic of chromatin fractionation analysis. 293T cells were transfected with mouse Aire expressing plasmids for 48 hrs before harvesting. Using a nuclear extraction buffer designed to preserve Aire-interaction partners^8^, we examined Aire solubility with or without micrococcal nuclease (MNase) treatment^9^. MNase treatment of nuclear extracts is known to solubilize chromatin along with proteins associated with chromatin^9^. Indeed, chromatin (inferred by levels of histone H3) exclusively partitions in the insoluble (P) fraction of nuclear extracts until the addition of MNase, which frees chromatin into the soluble (S) fraction. Similar to H3 partitioning, WT Aire remains mostly insoluble until chromatin is solubilized with MNase treatment, which releases a portion of Aire. A significant portion of WT Aire remained in the P fraction even after MNase digestion, possibly due to the formation of large Aire polymers that cannot be dissolved in our nuclear extraction buffer. In contrast to WT, chromatin fractionation of ΔPHD1 showed no interaction with chromatin. ΔPHD1 existed solely in the P fractions independent of MNase treatment, indicating the formation of insoluble aggregates that are not associated with chromatin. (D) Average H3K27ac and H3K4me3 ChIP profiles (normalized CPM) centered at WT-preferred sites versus ΔCTT.R3-preferred sites (top and bottom 500 peaks from Fig. 4A, respectively). Both H3K27ac and H3K4me3 ChIPs were performed in 4D6 cells prior to Aire expression.

