## Supplementary material for "Mechanism for controlled assembly of transcriptional condensates by Aire": Methods

***Expression vectors***

Throughout the Methods, Aire indicates human Aire, unless mentioned otherwise. Generation of pInducer20-Aire-FLAG WT and C311Y, pCDNA3.1-Aire-FLAG along with truncation variant aa 106-545 (Aire ΔCARD), pEGFP-N1-mouse Aire-FLAG (EGFP gene removed) WT along with truncation variant aa 108-552 (mouse Aire ΔCARD) and pEGFP-N1-mouse Aire aa 1-173 fused to monomeric enhanced GFP (mGFP) with a 3XFLAG-TEV protease cleavage site linker between the mouse Aire NLS and mGFP (CARD-mGFP) were previously described^1^. Lentiviral vectors pMD2.G (encoding VSV-G), psPAX2 (encoding HIV Gag/Gag-Pol, Rev, and Tat), and pInducer20-GFP (encoding both tetracycline repressor and GFP downstream of a tetracycline response element) were generous gifts from Dr. Hidde Ploegh (Harvard Medical School; Boston, MA). All point mutations in this study were generated by using Phusion® High Fidelity DNA polymerase (New England Biolabs). The same mutagenesis strategy was used to introduce a stop codon and FLAG tag after aa 480 to generate pEGFP-N1 mouse Aire-FLAG Δaa 480-552 (ΔCTT). To generate pInducer20-Aire-FLAG K83E and G228W, point mutations were first incorporated within pCDNA3.1-Aire-FLAG and then DNA encoding Aire-FLAG point mutants were subcloned into pInducer20-GFP (GFP gene removed). Generation of all internal amino acid deletions of Aire plasmids were generated by inverse PCR of template plasmid using PrimeSTAR Max DNA polymerase (Takara Bio), followed by DpnI (New England Biolabs) template digestion, 5’ phosphorylation and subsequent ligation of PCR products with T4 protein kinase and T4 DNA ligase, respectively (New England Biolabs). Aire Δaa 295-343 (ΔPHD1), Δaa 482-545 (ΔCTT), Δaa 499-509 (ΔCTT.R1), Δaa 510-521 (ΔCTT.R2), Δaa 522-535 (ΔCTT.R3) were first subcloned into pCDNA 3.1; subsequently, DNA encoding these Aire-FLAG variant constructs replaced DNA encoding GFP in pInducer20-GFP. DNA encoding APEX2 (cDNA derived from Addgene plasmid #49386^2^, a gift from Dr. Alice Ting) and Myc NLS-mouse Aire aa 486-552 was subcloned into pCDNA3.1. pGL4.31 (Firefly luciferase reporter plasmid under 5XUAS box promoter) was a kind gift from Dr. George Church (Harvard Medical School; Boston, MA). phRLCMV (Renilla luciferase expression plasmid under CMV promoter) was a kind gift from Dr. Diane Mathis (Harvard Medical School; Boston, MA). DNA encoding GAL4 DNA binding domain (DBD) was amplified from pMBD-Gate2, a kind gift from Kamil Onder (Addgene plasmid # 25947)^3^ and subcloned into pFLAG-CMV4 (pFLAG-CMV4-GAL4). mouse Aire aa 491-552 (CTT) WT, point mutants along with deletion variants Δaa 503-513 (ΔCTT.R1) Δaa 514-525 (ΔCTT.R2) and Δaa 526-539 (ΔCTT.R3) were subcloned into pFLAG-CMV4-GAL4. lentiCas9-Blast was a gift from Feng Zhang (Addgene plasmid # 52962)^4^. pLN426 (FuGW-G5p-mKate2) was a gift from Timothy Lu (Addgene plasmid # 105183)^5^. EGFP-P2A-Gal4DBD-mouse AireCTT was subcloned into pInducer20. FLAG tag, mouse Aire aa 486-552 (CTT construct used only for mass spectrometry) along with mouse Aire aa 503-552-FLAG (CTT-FLAG) WT, point mutants and deletion variants were subcloned into a modified pGEX-6P-1 vector containing a 6XHis-tag N-terminal to the GST tag. mouse Aire-FLAG point mutants and domain deletion variants Δaa 198-264 (ΔSAND), Δaa 301-342 (ΔPHD1), Δaa 433-475 (ΔPHD2), ΔCARDΔCTT, ΔSANDΔCTT, ΔPHD1ΔCTT, ΔPHD2ΔCTT, ΔSANDΔPHD2 (for clarity referred to as CARD-PHD1-CTT), ΔSANDΔPHD2ΔCTT (for clarity referred to as CARD-PHD1), and aa 1-290 (CARD-SAND) were subcloned into pEGFP-N1 (EGFP gene removed). pEGFP-N1 mouse Aire CARD-PHD2 was generated by replacing the DNA encoding for mGFP with DNA encoding mouse Aire PHD2 (aa 433-475) in pEGFP-N1 CARD-mGFP. pSG5-HA-p300 was a gift from Elizabeth Wilson (Addgene plasmid # 89094)^6^. Generation of HA-p300 truncation variants (amino acid boundaries listed in Figure S4) involved the same inverse PCR strategy as described above. For mGFP-p300 variant fusions, p300 variants were subcloned into a modified pEGFP-N1 vector where A206K point mutation was introduced in enhanced GFP to encode monomeric GFP. pET22b-HisGb1-STAT1(aa 710-750) + CBP TAZ2 (aa 1764-1855) was a gift from Peter Wright (Addgene plasmid # 99342)^7^. mouse Aire CTT (aa 480-552) and mouse Aire PHD1 (aa 295-349) were subcloned into pET47b. DNA encoding CHD4 PHD2 (aa 439-493) was synthesized by Integrated DNA Technologies (IDT); this cDNA was used as a template for subcloning into pET47b. cDNA for Sp110 CARD (aa 6-110) was amplified from MegaMan Human Transcriptome Library (Stratagene). The In-Fusion HD assembly method (Takara Bio) was used to generate CHD4-mouse Aire chimeras along with Sp110 CARD fusions with mGFP and mouse Aire PHDs within pEGFP-N1.

***Cell culture and transfection***

293T cells were maintained in DMEM supplemented with 10% FBS, 1% L-glutamine. 293T cells were transfected with either polyethyleneimine (PEI, 3.75 μg per well of 6-well plate with 1.5 µg DNA) or Lipofectamine 2000 (Invitrogen, 1 µg DNA per well of 12-well plate) according to manufacturer’s protocol. 4D6 cells were maintained in RPMI supplemented with 10% FBS, 1% L-glutamine, and transfected with Lipofectamine 2000 or Lipofectamine 3000 (Invitrogen) according to manufacturer’s protocol (1 µg DNA per well of 12-well plate).

For inhibitor treatments, 4D6 stable cells were induced with Dox for indicated amounts of time. 4 hrs prior to harvesting Dox-treated cells for different assays, equal volume of DMSO, A-485 (3 μM final concentration, Tocris Bioscience), dCBP-1 (0.25 μM final concentration, MedChemExpress), dBET6 (100 nM final concentration, MedChemExpress), or JQ1 (1 μM final concentration, Selleck Chemical) were added to medium.

For stable 4D6 cell line generation, lentivirus was first produced in 293T cells. 293T cells were seeded in a 12-well plate format and each well was transfected with 0.75 µg pInducer20, 0.33 µg psPAX2, and 0.18 µg pMD2.G using Lipofectamine 2000. 16 hours later, the medium was replaced. 48 hours after transfection, the medium (inoculum) was harvested and passed through a 0.45µm filter. For transduction, 4D6 cells were seeded in a 12-well plate. When cells were 70% confluent, the medium was replaced with a mixture of 300 µl filtered inoculum + 3 µl polybrene (10 mg/ml stock concentration, Sigma-Aldrich). Cells were incubated in the inoculum mixture with manual gentle agitation every 15 minutes for 1 hr. 700 µl of RPMI supplemented with 10% FBS, 1% L-glutamine was then directly added to each well with cells. 24 hours later, the transduced cells were trypsinized and transferred into a T25 flask to recover from transduction. After 24 hrs of recovery, 1 mg/ml G418 sulfate (Corning) was used for selection of transduced 4D6 cells. During G418 selection, medium was changed every 2-3 days. After the mock-transduced cells were ~95% dead, cells undergoing G418 selection were diluted into 96-well plates for individual clone selection. For each stable 4D6 clone, a doxycycline (Dox) titration curve was used to determine the appropriate Dox concentrations to use to have similar expression levels of Aire-FLAG variants compared to WT Aire-FLAG expression with 1 µg/ml Dox. 1, 0.1, 0.1, 0.1, 0.05, 1 µg/ml Dox was used on stable 4D6 clones expressing Aire-FLAG K83E, G228W, C311Y, ΔCARD, ΔPHD1, ΔCTT, respectively. 1, 1, and 0.1 µg/ml Dox was used on stable 4D6 clones expressing Aire-FLAG ΔCTT.R1-3, respectively. For ChIP-seq experiments with stable 4D6 cells treated with dCBP-1 for 4 hrs, 0.55 µg/ml Dox was used in order to induce the same expression levels of WT Aire as stable 4D6 cells treated with 1 µg/ml Dox + DMSO.

***Antibodies***

Antibodies used for immunofluorescence (IF) microscopy were mouse anti-FLAG (M2, Sigma-Aldrich, F1804), mouse anti-FLAG conjugated with FITC (M2, Sigma-Aldrich, F4049), rabbit anti-p300 (D8Z4E, Cell Signaling Technology, 86377S), rabbit anti-CBP (D6C5, Cell Signaling Technology, 7389S), rabbit anti-MED1 (Novus Biologicals, NB100-2574), Alexa Fluor® 488 AffiniPure donkey anti-mouse IgG (Jackson ImmunoResearch, 715-545-151), Alexa Fluor 647 AffiniPure donkey anti-rabbit IgG (Jackson ImmunoResearch, 711-605-152). Antibodies used for immunoblotting were rabbit anti-beta-actin (Cell Signaling Technology, 8457S), rabbit anti-HA (C29F4, Cell Signaling Technology, 3724S), mouse anti-FLAG-HRP (M2, Sigma-Aldrich, A8592), mouse anti-Lamin A (133A2, Cell Signaling Technology, 86846), mouse anti-Histone H3 (Cell Signaling Technology, 14269S), rabbit anti-Histone H3K27ac (D5E4, Cell Signaling Technology, 8173S), rabbit anti-Histone H3K18ac (D8Z5H, Cell Signaling Technology, 13998), rabbit anti-p300 (D8Z4E, Cell Signaling Technology, 86377S), rabbit anti-CBP (D6C5, Cell Signaling Technology, 7389S), rabbit anti-actyl-p300/CBP (Cell Signaling Technology, 4771S), anti-rabbit IgG-HRP (Cell Signaling Technology, 7074), anti-mouse IgG-HRP (Cell Signaling Technology, 7076). Antibodies used for chromatin immunoprecipitation mouse anti-FLAG (M2, Sigma-Aldrich, F1804), rabbit anti-Histone H3K27ac (D5E4, Cell Signaling Technology, 8173S), rabbit anti-Histone H3K4me1 (D1A9, Cell Signaling Technology, 5326S), rabbit anti-Histone H3K27me3 (C36B11, Cell Signaling Technology, 9733S), rabbit anti-Histone H3K4me0 (Active Motif, 91317), rabbit anti-p300 (D2X6N, Cell Signaling Technology, 54062), spike-in antibody (Active Motif, 61686).

***CRISPR/Cas9 screening and analysis***

The 4D6 cell line was transduced with lentiCas9-Blast (Addgene plasmid # 52962)^4^ and selected under 10 mg/ml blasticidin, and clones were picked for homogeneous Cas9 expression. Cas9-expressing 4D6 cells were transduced with FuGW-G5p-mKate2 (Addgene plasmid # 105183)^5^ and clones were picked and verified of UAS-mKate2 genomic insertion by PCR. PCR primers used for mKate2: forward CCTCACTCTAGATCTGCGATCT; reverse CGACCACCTTGATTCTCATGGT. PCR primers used for Cas9 as a control: forward GTTTGCCGCCAGAACACAGGA; reverse CACCTTGTACTCGTCGGTGATCA. Then, the Cas9+UASmKate2-expressing cells were transduced with pInducer20-EGFP-P2A-Gal4DBD-mouse AireCTT and clones with homogenous EGFP expression after 1 µg/ml Dox treatment were picked and further verified of AireCTT-dependent mKate2 expression by expressing sgRNAs that target mouse AireCTT. sgRNAs used to target mouse AireCTT: #1 TCCAGCACCTGGGCTTGCCA; #2 AGGTGCTGGACGGGGCCCAG. sgRNA that targets VP16 activation domain was used as a negative control: CCCCGACCGATGTCAGCCTG.

The engineered 4D6 cells were transduced with lentiviral Human Brunello CRISPR knockout pooled sgRNA library [a gift from David Root and John Doench (Addgene #73178-LV)^8^] at a MOI of 0.4, aiming for 500-fold representation of each sgRNA. Library transduced cells were selected under 1 mg/ml puromycin for 2 days and further expanded for another 7 or 10 days. Cells were treated with 1 µg/ml Dox 24 hrs prior to sorting to induce EGFP-P2A-Gal4DBD-AireCTT expression, and then top-5% and bottom-5% of the population were sorted based on mKate2/EGFP ratios on the SH800S Cell Sorter (Sony Biotechnology). Genomic DNA was extracted using the DNeasy Blood and Tissue Kit (QIAGEN, #69504) and cleaned using the OneStep PCR Inhibitor Removal Kit (Zymo Research, #D6030). Sequencing libraries were generated by PCR amplification as previously described^8^, pooled at an equal molar concentration, purified using the MinElute Reaction Cleanup Kit (Qiagen, #28204) to enrich for the 350-360bp amplicons, and subsequently sequenced on an Illumina sequencing platform (GENEWIZ). Demultiplexed sequencing reads were trimmed using Cutadapt (v2.5)^9^ to remove vector-derived sequences, yielding only 20 bp sequences corresponding to sgRNAs. The statistical analysis of sgRNA enrichment (Supplementary Table 3) was performed using MAGeCK-VISPR (v0.5.6) with the "MAGeCK-RRA" experimental configuration and visualized using the MAGeCKFlute R package (v2.0.0)^10^.

***RNA-seq and analysis***

Dox-inducible Aire-expressing 4D6 cells were cultured in the absence or presence of 1 µg/mL dox for 24 hrs. Total RNA was immediately extracted by using Direct-zol RNA Miniprep Kit (Zymo Research, R2052) with DNase I digestion. RNA-seq libraries were prepared using the NEBNext Ultra II RNA Library Prep Kit (New England Biolabs, E7775S) with ribosomal RNA depletion using the NEBNext rRNA Depletion Kit v2 (New England Biolabs, E7405L) and sequenced on the Illumina NovaSeq 6000 system (Novogene) with 150bp paired-end reads.

QC was performed on demultiplexed sequencing files using FASTQC (v0.11.3)^11^. Sequencing reads were trimmed using Trimmomatic (v0.36) and aligned to reference genome (GRCh38 primary assembly, release v43) using STAR (v2.7.0a)^12^. Read counting across genomic features was performed using featureCounts function within the Rsubread R package (v2.12.3) with duplicated reads ignored (ignorDup=T)^13^. Differential gene expression analysis was performed using the DEseq2 R package (v1.38.3) and visualized using the ggplot2 R package (v3.4.1)^14, 15^. BAM files generated during the STAR alignment were converted to bigwig files using deepTools (v3.5.1)^16^ with the following settings: bamCoverage --scaleFactor "scale factor" --smoothLength 150 --binSize 50 -e 200. "scale factor" was calculated as 1 divided by size factor that was obtained during DEseq2 analysis. After verification of consistency between replicates, bigwig files were averaged using WiggleTools (v1.2.2) and bedGraphToBigWig (v366) and imported into Integrative Genomics Viewer (IGV, v2.15.1) for visualization at specific loci^17, 18, 19^.

***5'-ethynyl uridine RNA-sequencing (EU-seq) and analysis***

Dox-inducible Aire-expressing 4D6 cells were cultured in the absence or presence of 1 µg/mL dox for 24 hrs. Equal volume of DMSO or 3 µM A-485 was added to cells 4 hrs prior, and 0.5 mM 5'-EU was added to the cell culture 30min prior to RNA extraction. Total RNA was immediately extracted by using Direct-zol RNA Miniprep Kit (Zymo Research, R2052) with DNase I digestion. Ribosomal RNA was depleted using the NEBNext rRNA Depletion Kit v2 (New England Biolabs, E7405L). 5'-EU labeled nascent RNA was biotinylated and pulled down using the Click-iT Nascent RNA Capture Kit (Invitrogen, #C10365) according to the manufacturer's protocol. 5'-EU labeled RNA captured on streptavidin-beads was then immediately used for sequencing library preparation using the NEBNext Ultra II RNA Library Prep Kit (New England Biolabs, E7775S). Libraries were sequenced on the Illumina NovaSeq 6000 system (Novogene) with 150bp paired-end reads.

QC was performed on demultiplexed sequencing files using FASTQC (v0.11.3). Sequencing reads were trimmed using Trimmomatic (v0.36) and aligned to reference genome (GRCh38 primary assembly, release v43) using STAR (v2.7.0a). BAM files generated during the STAR alignment were converted to bigwig files using deepTools (v3.5.1) with the following settings: bamCoverage --scaleFactor "scale factor" --smoothLength 150 --binSize 50 -e 200. "scale factor" was calculated as 1 divided by size factor that was obtained during DEseq2 analysis. After verification of consistency between replicates, bigwig files were averaged using WiggleTools (v1.2.2) and bedGraphToBigWig (v366). Bigwig files showing log_2_ fold-changes between two groups were generated using deepTools (v3.5.1) with the bigwigCompare function. Bigwigs files were imported into Integrative Genomics Viewer (IGV, v2.15.1) for visualization at specific loci.

***Chromatin immunoprecipitation (ChIP)-seq and analysis***

4D6 stable cells were seeded on 150mm plates with or without Dox and grown for 24 hrs. For drug-treated samples, equal volume of DMSO, 3 µM A-485, or 0.25 µM dCBP-1 was added to cells 4 hrs prior to harvest. For anti-FLAG and anti-p300 ChIP-seq, cells were washed 3 times with PBS and then crosslinked with 2mM Disuccinimidyl glutarate (DSG) in PBS for 45 minutes at room temperature. Cells were then washed again 3 times with PBS and then crosslinked with 1% formaldehyde (Sigma, Thermofisher, and Electron Microscopy Sciences) in PBS for 10 minutes at room temperature. For anti-histone mark ChIP-seq, cells were crosslinked with 1% formaldehyde in fresh media for 10 minutes at room temperature. After formaldehyde crosslinking, all cells were washed 1 time with PBS and quenched with 0.125 M glycine in PBS for 5 min at room temperature. Quenched cells were washed with ice cold PBS, then harvested in ice cold PBS supplemented with 0.5 mM PMSF. Cells were spun down at 500 g for 5 minutes and cell pellets were supplemented with 1 µl of 100mM PMSF and 1 µl 1X mammalian protease inhibitor cocktail (G-Biosciences), then flash frozen in liquid nitrogen and stored at -80°C until ready to use.

For one ChIP pull-down, ~15x10^6^ cells were used. Cells pellets were thawed on ice for 15 minutes and then resuspended in ice-cold One-Step Lysis Buffer (50 mM Tris H 7.5, 1% SDS, 0.25% Sodium Deoxycholate and 1 X mammalian protease inhibitor cocktail). Cells were incubated on ice for 10 min and then 700 µl ChIP Dilution Buffer (50 mM Tris pH 7.5, 0.01% SDS, 150 mM NaCl, 0.25% Triton-X, 1 X mammalian protease inhibitor cocktail) was added. Chromatin was sheared using with a Covaris M220 ultrasonicator (settings: 5% duty factor, 75W max power, 200 cycles per burst, 20 min) at 6°C. Lysates were spun down in a refrigerated centrifuge for 10 minutes at 18,000 g. Cleared lysate was 3-fold diluted with ChIP Dilution Buffer, spike-in chromatin (600ng per pull-down, Active Motif) was added when cells were treated with p300 inhibitors or DMSO, and 2-3% of the input was saved for later use. Lysates were nutated with antibodies [for each pull-down when indicated, 2.5 µg of anti-FLAG (M2, Sigma); 5 µg anti-H3K4me1, 5 µg anti-H3K27me3, 4 µg of anti-p300 (D2X6N, Cell Signaling Technologies); 4 µg of anti-H3K27Ac (Cell Signaling Technologies); 4 µg of anti-H3K4me0 (Active Motif); an additional 2 µg spike-in antibody (Active Motif) was included when cells had been treated with p300 inhibitors or DMSO] for 16 hrs at 4°C.

Protein-DNA complexes were immunoprecipitated using Protein G magnetic beads (Active Motif) with 2 hrs nutation at 4°C. Protein G beads were washed with the following ice-cold buffers: RIPA buffer (0.1% SDS, 0.1% Sodium Deoxycholate, 1% Triton X-100, 1 mM EDTA, 10 mM Tris pH 8, 150 mM NaCl); RIPA supplemented with 350 mM NaCl; LiCl Buffer (10 mM Tris pH 8, 250 mM LiCl, 0.5% Triton X-100, 0.5% Sodium Deoxycholate) and Tris Buffer (10 mM Tris pH 8.5). Protein-DNA complexes were eluted with Elution Buffer (10 mM Tris pH 8, 1 mM EDTA, 0.1% SDS, 150 mM NaCl, 5 mM DTT) with gentle agitation at 65°C for 1 hr. Reserved ChIP inputs were diluted two-fold with Elution Buffer. Dilute inputs and eluted protein-DNA were treated with RNAse (Machery-Nagel) at 37°C for 30 min, then Proteinase K (New England Biolabs) at 65°C for 16 hrs to ensure reverse crosslinking of DNA. Reverse cross-linked DNA was purified using SPRI Select beads (Beckman). ChIP-seq DNA libraries were prepared using NEBNext Ultra II DNA library Prep Kit for Illumina (New England Biolabs) according to the manufacturer’s protocol. Deep sequencing was performed using a NovaSeq sequencer (Illumina) with pair-end 150 bp reads.

QC was performed on demultiplexed sequencing files using FASTQC (v0.11.3). Sequencing reads were trimmed using Trimmomatic (v0.36) and aligned to reference genome (GRCh38 primary assembly) using bwa (v0.7.17). The resulting SAM files were converted to BAM files, sorted and indexed using Samtools (v1.6). Samples with drosophila spike-in chromatins were aligned to a customized reference genome consisting both hg38 and dm6, and the sorted and index BAM files were subsequently split into separate BAM files containing hg38- and dm6-mapped reads, respectively, using plit_bam.py module in the SPIKER tool^20^. hg38- and dm6-mapped reads were deduplicated using the alignmentSieve function in deepTools (v3.5.1), and unique dm6-mapped reads were counted to calculate the scaling factors. Peak calling was performed using MACS2 (v2.2.7.1)^21^ with the following parameters: macs2 callpeak -f BAMPE -B -g 3.2e+9 --keep-dup 1 --SPMR --nomodel --extsize 250 -q 0.05 --cutoff-analysis. When calling peaks in WT and mutant Aire ChIP-seq samples, corresponding input controls were included to remove the effect of background noise. WT and mutant Aire ChIP-seq BAM files and peaks were imported into DiffBind (v3.8.4)^22^, and a census of all Aire peaks (n=13023, summits +/– 200 bp, Supplementary Table 5) were obtained after removing blacklisted and greylisted regions using dba.blacklist and dba.peakset functions. Trimmed Mean of M-values (TMM)-normalized counts-per-million (CPM) reads^23^ within each Aire peak (summits +/– 200 bp) were obtained using the dba.count function in DiffBind with options: bRemoveDuplicates = T, score=DBA_SCORE_TMM_MINUS_FULL_CPM. TMM-normalized CPM reads of H3K27ac ChIP and p300 ChIP around Aire peaks (Aire peak summits +/­– 2000 bp for H3K27ac and summits +/– 200 bp for p300) were obtained with option score=DBA_SCORE_TMM_READS_FULL_CPM. Statistical comparison and significance were calculated using the Wilcoxon rank sum test in R. Spearman's correlation coefficient was calculated in Prism (Graphpad). Peaks called from H3K27ac ChIP-seq (­–Aire expression) were removed of blacklisted regions and subjected to Ranking Ordering of Super Enhancer (ROSE) analysis^24^ to obtain putative super-enhancer regions, using 12.5 Kb stitching distance and 2 Kb TSS exclusion parameters. BAM files were converted to bigwig files using deepTools (v3.5.1). Bigwig files for WT and mutant Aire ChIP-seq samples shown in Figures 1D, 4A, 4C, 6D, S1A, S1F were generated by subtracting background noises in corresponding input controls with the following settings: bamCompare -b1 "pulldown bam file" -b2 "input bam file" -o "bw file" --operation subtract --scaleFactorsMethod None --normalizeUsing CPM --ignoreDuplicates -bs 50 -e 200 --smoothLength 150. Bigwig files for all other sequencing samples were generated without input controls using the bamCoverage function with the same setting options. After verification of consistency between replicates, bigwig files were averaged using WiggleTools (v1.2.2) and bedGraphToBigWig (v366) and imported into IGV (v2.15.1) for visualization at specific loci or into deepTools (v3.5.1) for generation of heatmaps and average profiles using the computeMatrix, plotHeatmap and plotProfile functions. The heatmaps shown in Figures 4A and 6D were centered on the census of all Aire ChIP-seq peak-defined regions (n=13023, +/– 2 Kb) and ranked based on the ratio of signals in WT versus DCTT.R3 samples. The heatmaps shown in Figures S1A, 4F-4G were centered and scaled at the H3K27ac-delimited super-enhancer regions (n=529, +/– 200 Kb) and ranked based on the ROSE ranking order.

***Transposase-accessible chromatin with sequencing (ATAC-seq) and analysis***

Stable 4D6 cells –/+ Dox were harvested 24 hrs after induction. 100,000 cells were used for each replicate. ATAC-seq libraries were prepared using ATAC-seq library prep kit (Active Motif) according to the manufacturer’s protocol. Deep sequencing was performed using a NovaSeq sequencer (Illumina) with pair-end 150 bp reads.

QC was performed on demultiplexed sequencing files using FASTQC (v0.11.3). Sequencing reads were trimmed using Trimmomatic (v0.36) and aligned to reference genome (GRCh38 primary assembly) using bwa (v0.7.17). The resulting SAM files were converted to BAM files, sorted and indexed, and reads mapped to mitochondrial DNA were removed using Samtools (v1.6). Read fragment sizes were checked using the ATACseqQC R package (v3.19)^25^. Post PCR duplicate removal, reads were shifted +4bp and –5bp for positive and negative strand respectively, and were further split into nucleosome-free regions (NFR), mono- or di-nucleosome regions using the alignmentSieve function in deepTools (v3.5.1). Peak calling was performed on bam files containing reads mapped to NFR using MACS2 (v2.2.7.1). NFRs that overlapped with Aire peaks and showed strong Aire-ChIP signals were selected as Aire-bound NFRs (n=542, Supplementary Table 1); whereas NFRs that had similar ATAC-seq read pileups as those in Aire-bound NFRs but showed no Aire-ChIP signals were selected as Aire-free NFRs (n=658, Supplementary Table 1).

***Immunofluorescence microscopy***

4D6 or 293T cells were seeded onto glass cover slips in 12-well plate format. Cells at ~70% confluence were transiently transfected with indicated plasmids. Dox-inducible Aire-expressing 4D6 cells were seeded in the presence or absence of Dox. For inhibitor treatments, 4D6 stable cells were induced with Dox for indicated amounts of time. 4 hrs prior to fixing Dox-treated cells, equal volume of DMSO, A-485 (3 μM final concentration) or dCBP-1 (0.25 μM final concentration), dBET6 (100 nM final concentration), or JQ1 (1 μM final concentration) were added to medium. 24 hrs post-transfection or Dox treatment for indicated amount of time, cells were washed with PBS, then fixed with 2% paraformaldehyde in PBS for 10 minutes. Cells were washed again with PBS and then permeabilized with 0.5% Triton X-100 in PBS for 10 minutes. Cells were blocked with 1% BSA in PBST (PBS + 0.2% Tween-20) for 15 minutes at room temperature or 16 hrs at 4°C, and then probed with antibodies. Cells were then counterstained with DAPI (Life Technologies). Coverslips were mounted using Fluoromount-G (SouthernBiotech) or Vectashield (Vector Laboratories, #H-1000-10) and then imaged on fluorescence microscopes. 2D images were captured on a wide-field Zeiss Axio Imager M1. Image z-stacks were captured on a wide-field Nikon Ti2 equipped with a Nikon DS-Qi2 large-format CMOS camera (11 frames per stack, 0.3 μm z-step), or a Yokogawa spinning disk confocal Nikon Ti equipped with a Hamamatsu ORCA-Fusion BT sCMOS camera (9 frames per stack, 0.3 μm z-step).

All quantitative immunofluorescence imaging analyses were performed with Fiji Is Just ImageJ (FIJI, ImageJ2 v2.14.0/1.54f). For each image z-stack, masks were drawn around nuclei based on DAPI fluorescence. Only cells that expressed Aire (anti-FLAG immunostaining) as visualized by nuclear fluorescence staining were analyzed further. For the calculation of mean fluorescence intensity per segmented nucleus, background subtraction was performed on each image to compare intensities between samples. To examine the number of Aire foci per nucleus, Aire foci were segmented using the Yen Dark thresholding method^26^ featured in FIJI. For Aire foci volume analyses of 4D6 cells, the DiAna ImageJ plug-in^27^ was used to 3D segment Aire foci and calculate Aire foci volumes using the same intensity thresholds when comparing samples.

To identify nuclei that contained Aire foci or “diffuse” Aire, image z-stacks were first manually inspected for the most in-focus 2D z-slice to further analyze. Within 2D images, a nucleus would be defined to have Aire foci with the presence of 2 or more circular spots (> 3 pixels^2^) with maximum intensities >2 fold higher than background intensities within the nucleus. Diffuse nuclear staining was defined as uniform Aire fluorescence intensities throughout a given nucleus.

For image analyses of stable 4D6 cells expressing Aire-FLAG WT or ΔPHD1 co-stained with co-activators, image z-stacks were first manually inspected for the most in-focus 2D z-slice to further analyze. To identify Aire foci in a 2D image, an iterative implementation of Minimum Error intensity thresholding was used to create region of interests (ROIs) outlining Aire foci for a given nucleus. For every identified Aire focus, the average intensity signal from the corresponding ROI in the co-activator fluorescence channel was determined. The average intensities were normalized to the average co-activator intensity within the entire nucleus being examined. Statistical significance comparisons were calculated by using a Mann-Whitney test for two population proportions where each population consists of all individually normalized mean intensities of co-activators within the corresponding locations of Aire foci.

***Immunofluorescence with RNA fluorescence in situ hybridization (FISH)***

Cells were plated on cover slips in 12-well tissue culture plates and grew for a total of 24 hrs. Dox and/or drugs were supplemented to media for the indicated times prior to fixation of cells. Cells were then washed with PBS once and fixed using 4% paraformaldehyde (Electron Microscopy Sciences, #15714) in PBS for 10 min. After washing cells twice in PBS, permeabilization of cells was performed using 0.5% Triton-X100 in PBS for 10 min, followed by washing with PBS-T twice. Cells were blocked with 1% RNase-free BSA (Sigma-Aldrich, #126609) in PBS-T for 30 min, and then incubated with the FITC conjugated anti-FLAG antibody (Sigma, F4049) at a final concentration of 10 μg/ml in PBS-T with 1% BSA at room temperature for 1 hr. After washing in PBS-T twice and PBS once, cells were re-fixed using 4% PFA in PBS for 10 min. After two washes of PBS, cells were pre-incubated in Buffer A [20% Stellaris RNA-FISH buffer A (Biosearch Technologies, #SMF-WA1-60) and 10% deionized formamide (Millipore, #S4117) in RNase-free water (Thermofisher, #10977023)] for 5 min. Cells were then incubated with a final concentration of 125 nM nascent RNA probes in hybridization buffer [90% Stellaris RNA-FISH hybridization buffer (Biosearch Technologies, #SMF HB1-10) and 10% deionized formamide] overnight in a humidified chamber at 37˚C. After washing with Buffer A for 30min at 37˚C, nuclei were stained with DAPI in Buffer A for 5 min, followed by a wash in Stellaris RNA-FISH buffer B (Biosearch Technologies, # SMF-WB1-20) for 5 min. Coverslips were mounted onto glass slides with Vectashield (Vector Laboratories, #H-1000-10) and sealed with nail polish. Nascent RNA-FISH probes were custom-designed to target *RIC8A*, *SETD1B* and *UBTF* intronic regions using the Stellaris probe designer and manufactured at Biosearch Technologies. The sequences of RNA-FISH probes are listed in Supplementary Table 2.

3D images were acquired on the inverted Nikon Ti2 fluorescence microscope with 60X objective using NIS-Elements acquisition software, at a resolution of 9.2308 pixels/mm and voxel size of 0.1083 x 0.1083 x 0.3 mm^3^. Microscope specifications can be found at https://nic.med.harvard.edu/microscopes/george_michael/. Images were post-processed using FIJI for further analyses.

Nascent RNA-FISH spots and Aire foci were identified in individual z-stacks using DiAna ImageJ plug-in^27^. For images generated from the same experiment, the "threshold" parameter for a given fluorescent channel was set the same across different groups, and the "min pixel" parameter was set to 20 for RNA-FISH spots and 5 for Aire foci. The distances between Aire foci and RNA-FISH spots or randomized nuclear spots were then measured using DiAna ImageJ plug-in^27^. Statistical comparison and significance were calculated using the Kolmogorov-Smirnov test in Prism (Graphpad).

The average Aire signal at RNA-FISH spots was computed and plotted in MATLAB as previously described^28^. Briefly, the MATLAB scripts were obtained from github (https://github.com/krishna-shrinivas/FISH_IF_colocalization), and a list of RNA FISH signal centroids (x, y, z) manually curated using DiAna from the previous step were provided to the MATLAB pipeline. The Aire IF signal centered at FISH spots were then combined to calculate an average intensity projection within a 2.8 x 2.8 mm^2^ square. The same process was carried out for FISH signal centered on its own (x, y, z) coordinates. Spearman correlation coefficient (*r*_s_) was computed and reported between the FISH and IF signals centered at FISH spots. As a control, the same process was carried out for Aire IF signal centered at random nuclear positions that were selected using this MATLAB pipeline. The average intensity projections were then used to generate 2D contour plots of the signal intensity. The averaged IF signal centered at FISH spots or randomly selected nuclear positions were plotted using the same color and intensity scale.

***Aire protein expression and chromatin fractionation assays***

4D6 and 293T cells were transfected with plasmids expressing indicated proteins for assaying expression levels and chromatin fractionation assays in 12-well and 6-well plate formats, respectively. Stable 4D6 cells induced with 0.1-1µg/ml Dox were also seeded in 12-well format for determining expression levels. 24 hours after transfection or doxycycline induction, cells were harvested in PBS and washed one more time with PBS. For samples expressing mouse Aire that were sensitive to protein degradation and/or histone deacetylation, washed cells were immediately lysed in 1% SDS Buffer (50 mM Tris pH 7.5, 150 mM NaCl, 1% SDS, 0.3 mM DTT; 75-100 µl/sample), boiled for 5 minutes and processed for western analyses as described below. For all other samples, washed cells were incubated in Hypotonic Buffer [20 mM HEPES pH 7.5, 0.05% IGEPAL, 1.5 mM MgCl_2_, 10 mM KCl, 5 mM EDTA, and 1X mammalian protease inhibitor cocktail; 50 µl and 100µl/sample for 12-well and 6-well plate formats, respectively] for 15 minutes at 4°C. The lysed cells were spun down at 500 g for 5 minutes at 4°C and the supernatant (cytoplasmic fraction) was removed. The pellet (nuclear fraction) was washed 2 times with ice-cold PBS.

For the comparison of mouse Aire variant expression levels, the PBS-washed nuclear fraction was lysed in 1% SDS Buffer and boiled for 5 minutes. BCA assay was used to determine the total protein concentration of lysates. Equal amounts of total protein from nuclear lysates were loaded on SDS-PAGE gel and subsequently analyzed by western blotting.

For the Aire chromatin fractionation assay, 293T cells transfected with plasmids expressing mouse Aire were harvested and washed in PBS, then lysed with Hypotonic Buffer as described above. Nuclear fractions were then resuspended in Nuclear Extraction Buffer (50 mM Bis-Tris pH 7.5, 750 mM 6-aminocaproic acid, 3 mM CaCl_2_, 10% glycerol, 1X mammalian protease inhibitor cocktail; 200 µl/sample) and split into fractions with or without MNase (Promega, 50U/100 µl of nuclear lysate) and incubated for 1 hour at 4°C. MNase activity was quenched with 5 mM EDTA. Nuclear lysates were centrifuged at 18,000 g for 10 minutes at 4°C. The resulting supernatant was the “soluble” nuclear fraction and saved for analysis. The “insoluble” nuclear pellet was washed one time with ice-cold PBS, then resuspended in Laemmli sample buffer and boiled for 5 minutes. The soluble and insoluble nuclear fractions were run on SDS-PAGE gel and subsequently analyzed by western blotting.

***Luciferase reporter assay***

4D6 cells were seeded into 48-well plates. At ∼80% confluence, cells were transfected with 200 ng pGL4.31 (Firefly luciferase reporter plasmid under 5XUAS box promoter), 1 ng phRLCMV (a constitutively expressed Renilla luciferase reporter plasmid) and 25 ng plasmid expressing Gal4-DBD-fusion variants by using Lipofectamine 2000 (Life Technologies) according to the manufacturer’s protocol. 24 hrs post-transfection, cells were lysed and the transcriptional activity of Gal4^DBD^-CTT was measured by using the Dual Luciferase Reporter assay (Promega) and a Synergy2 plate reader (BioTek). Firefly luciferase activity was normalized against Renilla luciferase activity.

***Quantitative real time PCR (RT-qPCR) and 5'-ethynyl uridine (EU)-qPCR***

4D6 stable cell lines expressing Dox-inducible WT or mutants were harvested for RNA extraction 24 hr post Dox treatment, and 4D6 cells transiently expressing mouse Aire were harvested for RNA extraction 24 hr post-transfection. For RT-qPCR, total RNA was immediately extracted using the Direct-zol RNA Miniprep Kit (Zymo Research, R2052) with DNase I digestion and reverse-transcribed using SuperScript II (Life Technologies) with oligo(dT_18_). For EU-qPCR, 0.5 mM 5'-EU was added to the cell culture 30min prior to RNA extraction. Total RNA was extracted the same way described above. 5'-EU labeled RNA was biotinylated and pulled down using the Click-iT Nascent RNA Capture Kit (Invitrogen, #C10365) according to the manufacturer's protocol. 5'-EU labeled RNA captured on streptavidin-beads was then immediately reversed transcribed using SuperScript II (Life Technologies) with oligo(dT_18_).

qPCR was performed using Power SYBR Green PCR Master Mix (Invitrogen) on a CFX-Connect detection system (Bio-Rad, Hercules, CA). The expression of Aire-induced genes was normalized against that of the Aire-independent gene *RPL18* using the ∆∆Ct method. *RPL31* (Aire-independent gene control) was also normalized against *RPL18*. The qPCR primer sequences are listed in Supplementary Table 2.

***E. coli. expression and purification of recombinant proteins***

His_6_-GST, His_6_-GST-FLAG, His_6_-GST-CTT, His_6_-GST-CTT-FLAG WT and variants were expressed in BL21(DE3) Rosetta (Millipore Sigma) cells at 37°C in Luria Broth (LB), grown to an OD_600_ ~0.8-1, then induced with 0.4 mM IPTG for 3 hrs at 37°C. His_6_-mouse Aire PHD1 and His_6_-CHD4 PHD2 were expressed in BL21(DE3) Rosetta cells at 37°C in LB, grown to an OD_600_ ~0.5-0.6, then cooled down to 25°C for 40-60 minutes while still shaking, supplemented with 50 μm ZnCl_2_ and induced with 0.4 mM IPTG for 5-6 hrs. His_6_-STAT1(710-750) + CBP TAZ2 (1764-1855) were expressed in BL21(DE3) Rosetta cells at 37°C in LB, grown to an OD_600_ ~0.8-1, then cooled down to 15°C for 15-20 min minutes on ice, supplemented with 150 μm ZnCl_2_ and induced with 0.4 mM IPTG for 16 hrs. His_6_-mouse Aire CTT-FLAG was co-expressed with pCDF-GroEL/ES+trigger factor (a generous gift from Timothy A. Springer lab, Boston Children’s Hospital; Boston, MA) in BL21(DE3) cells at 37°C in M9 minimal medium supplemented with ^15^NH_4_Cl and ^13^C_6_-glucose; cells were grown to OD_600_ ~0.8 and cooled down to 25°C, then induced with 0.5 mM IPTG for 5 hrs.

Cells expressing His_6_-GST and His_6_-GST-CTT for mass spectrometry were harvested and resuspended in MS Lysis Buffer (50 mM Tris pH 8, 300 mM NaCl and 10% glycerol, 1 mM PMSF). Cells expressing His_6_-GST-FLAG, His_6_-GST-CTT-FLAG variants, His_6_-mouse Aire PHD1, His_6_-CHD4 PHD2 and ^15^N^13^C-labeled His_6_-mouse Aire CTT were harvested and resuspended in FP lysis buffer (50 mM HEPES pH 7.5, 200 mM NaCl, 5% glycerol, 1 mM PMSF). Cells expressing STAT1(710-750) + CBP TAZ2 (1764-1855) were resuspended in TAZ2 lysis buffer (50 mM Tris pH 8, 400 mM NaCl, 10 mM MgCl_2_, 1 mM PMSF) supplemented with 10mM imidazole. All resuspended cell pellets were frozen and stored at -20°C until ready for purification. All protein purification procedures were performed at 4°C. All thawed cells were lysed with an Emulsiflex C3 (Avestin) and centrifuged at 32,000 g for 30 minutes. For each protein, cleared lysate was loaded onto a Ni^2+^-NTA agarose (Qiagen) gravity-flow column.

For purification of His_6_-GST and His_6_-GST-CTT, the Ni^2+^-NTA agarose columns were washed with 100 column volumes of MS Wash Buffer (50 mM Tris pH 8, 300 mM NaCl, 25 mM imidazole) and purified protein was eluted with 50 mM Tris pH 8, 300 mM NaCl and 50 mM - 250 mM imidazole. Imidazole elutions containing >95% pure protein were pooled and buffer exchanged back into MS Lysis Buffer using Amicon Ultra - 15 concentrators (Millipore) until ready for further use.

For purification of His_6_-GST-FLAG, His_6_-GST-CTT-FLAG variants, His_6_-mouse Aire PHD1 and His_6_-CHD4 PHD2, and ^15^N^13^C-labeled His_6_-mouse Aire CTT, the Ni^2+^-NTA agarose columns were washed with 7 column volumes of FP Wash Buffer (50 mM HEPES pH 7.5, 300 mM NaCl, 25mM Imidazole). Purified protein was eluted with 50 mM HEPES pH 7.5, 300 mM NaCl and 50 mM - 150 mM imidazole. Imidazole elutions containing >90% pure protein were pooled and supplemented with 5 mM BME. His_6_-GST-FLAG, His_6_-GST-CTT-FLAG variants, and His_6_-CHD4 PHD2 were dialyzed in FP Dialysis Buffer (25 mM HEPES pH 7.5, 200 mM NaCl, 5 mM BME) for 16 hrs. Dialyzed His_6_-GST-FLAG and His_6_-GST-CTT-FLAG variants were further purified with a Superdex 200 Increase 10/300 GL column (Cytiva) with SEC Buffer (25 mM HEPES pH 7.5, 100 mM NaCl, 5 mM BME). Dialyzed His_6_-CHD4 PHD2 was buffered exchanged into HEPES Loading Buffer (25 mM HEPES pH 8, 25 mM NaCl, 5 mM BME), then loaded onto a HiTrap Q FF 1 ml column (Cytiva) and purified with a gradient (Buffer A: 25 mM HEPES pH 8, 5 mM BME; Buffer B: 25 mM HEPES pH 8, 1 M NaCl, 5 mM BME). Dialyzed ^15^N^13^C-labeled His_6_-Aire CTT was buffered exchanged into Tris Loading Buffer (25 mM Tris pH 8, 25 mM NaCl, 5 mM BME), then loaded onto a Resource Q 1 ml column (Cytiva) and purified with a gradient (Buffer A: 25 mM Tris pH8, 5 mM BME; Buffer B: 25 mM Tris pH 8, 1 M NaCl, 5 mM BME). Ion-exchange pooled fractions of His_6_-CHD4-PHD2 and ^15^N^13^C-labeled His_6_-Aire CTT, and His_6_-Aire PHD1 were further purified with a Superdex 75 Increase 10/300 GL column (Cytiva) with SEC Buffer. Purified His_6_-GST-FLAG, His_6_-GST-CTT-FLAG variants, His_6_-Aire PHD1, and His_6_-CHD4 PHD2 were concentrated using Vivaspin 2 concentrators (Sartorius), flash frozen in liquid nitrogen and stored at -80°C until ready for further use. ^15^N^13^C-labeled His_6_-Aire CTT was concentrated and dialyzed into NMR Sample Buffer (10 mM NaPO_4_ pH 6.3, 30 mM NaCl, 1 mM DTT).

For His_6_-STAT1(710-750) + CBP TAZ2 (1764-1855), cleared lysate and Ni^2+^-NTA agarose beads were nutated for 2 hrs. Beads were then washed with 10 column volumes of the following: TAZ2 lysis buffer supplemented with 10 mM Imidazole; TAZ2 lysis buffer supplemented with 40 mM Imidazole; 50 mM Tris pH 8, 1 M NaCl; and 50 mM Tris pH 7.5, 500 mM NaSO4. His_6-_STAT:CBP complex was eluted with 50 mM Tris pH 7.5, 600 mM Imidazole. was buffered exchanged into MES Loading Buffer (25 mM MES pH 7, 50 mM NaCl, 5 mM BME), then loaded onto a Resource S 1 ml column (Cytiva) and purified with a gradient (Buffer A: 25 mM MES pH 7, 5 mM BME; Buffer B: 25 mM MES pH 7, 1 M NaCl, 5 mM BME). Ion-exchanged fractions containing only CBP TAZ2 were pooled, concentrated, flash frozen in liquid nitrogen and stored at -80°C until ready to use for GST-CTT-FLAG pull-downs; alternatively, pooled fractions were directly dialyzed into NMR Sample Buffer.

***Mass spectrometry of Aire CTT binding partners***

Equal amounts of His_6_-GST and His_6_-GST-mouse Aire CTT in MS Lysis Buffer were captured onto glutathione Sepharose beads (Cytiva) for 1 hr at 4°C. GST-protein bound beads were washed 3 times with MS Lysis Buffer. 293T cells were harvested and lysed in Hypotonic Buffer and Nuclear Extraction buffer to obtain nuclear “soluble” extracts as described above for chromatin fractionation assays. The 293T nuclear extracts were incubated with GST-protein bound beads for 16 hrs at 4°C. Beads were washed 1 time with ice-cold PBS supplemented with 0.05% IGEPAL and 3 times with ice-cold PBS. Bound proteins were eluted with Laemmli sample buffer and boiled for 5 minutes. The eluted proteins were run on SDS-PAGE gel and stained with Coomassie Blue. A band around ~250 kDa was cut out of the gel for each lane containing either proteins interacting with His_6_-GST-CTT or His_6_-GST as a negative control. Extracted gel bands were sent to the Taplin Mass Spectrometry Facility at Harvard Medical School for in-gel protease digestion and micro-capillary LC/MS/MS analysis for protein binding partner identification. Protein candidates identified from Mass Spectrometry were ranked from high to low based on their coverages (%) in His_6_-GST-CTT pulldown samples where there were no coverages in His_6_-GST pulldown samples (Supplementary Table 4).

***GST-Aire CTT-FLAG pull downs***

All pull-downs were performed at 4°C unless otherwise specified. Equal amounts of His_6_-GST-FLAG and His_6_-GST-mouse Aire CTT-FLAG variants in SEC Buffer were captured onto glutathione Sepharose beads for 30 min. GST-protein bound beads were washed 3 times with SEC Buffer, then blocked with 0.1% BSA in SEC Buffer for 30 min until ready to mix with 293T nuclear extracts or purified recombinant proteins.

For pull-downs with 293T lysates, 293T cells were transiently transfected with HA-p300 variant expression vectors in a 6-well plate format. 48 hours after transfection, cells were harvested and lysed in Hypotonic Buffer and Nuclear Extraction buffer to obtain nuclear “soluble” extracts as described above for chromatin fractionation assays. The nuclear fractions were directly mixed with BSA-blocked GST-protein-bound beads for 1 hr. For pull-downs with recombinant FLAG-tagged full-length p300 (Active Motif) was diluted into 25 mM HEPES pH 7.5, 100 mM NaCl, 1.5 mM MgCl_2_, 5 mM BME and added to BSA-blocked GST-protein-bound beads for 1 hr. Pull-downs with 293T lysates and recombinant full-length p300 were eluted by boiling beads in Laemmli sample buffer for 5 minutes.

For pull-downs with recombinant CBP TAZ2 protein, CBP TAZ2 was mixed with BSA-blocked GST-protein-bound beads in SEC Buffer for 1 hr, then beads were washed 3 times with SEC Buffer. Bound CBP TAZ2 was eluted with the addition of 25 mM glutathione (Sigma) in SEC Buffer. Elutions were passed through a Costar Spin-X centrifuge tube 0.22μm filter (Millipore) to prevent contaminating beads from entering samples.

For all pull-downs, input samples and pulled-down proteins were run on SDS-PAGE gel and analyzed by Krypton staining or western blotting.

***NMR spectroscopy***

A 15N T2 relaxation experiment was acquired at 15°C on a 700 MHz spectrometer, equipped with a TXI probe. The experiment was recorded as a pseudo3D in an interleaved manned using the standard Bruker pulse sequence “hsqct2etf3gpsi3d”. Relaxation delays of 16.3, 32.6, 65.2, 130.4, 163, 195.6, 228.2 and 260.8 ms were used; 32.6 ms was measured twice for error evaluation. An interscan delay of 5 sec was used and 20 scans were collected for each increment. 128 points were collected in the indirect ^15^N dimension with a spectral width of 22 ppm, 1024 point were collected in the direct ^1^H dimension with a spectral width of 16.2 ppm. The carrier was centered at 4.7 ppm in the ^1^H dimension and at 117 ppm in the indirect ^15^N dimension. Spectra were processed with NMRPipe^29^ and the decay fitted CCPNmr^30^. Data analysis was performed excluding overlapping resonances.

Spectra were also recorded on an 800MHz Bruker Advance spectrometer equipped with a TCI cryoprobe with z-shielded gradients and an Avance III console at 15°C. 3D-experiments were performed using NUS, collecting 10% of the Nyquist grid in the indirect dimension, using Poisson-Gap sampling. The resulting non-uniformly sampled spectra were reconstructed using the hmsIST algorithm^31^. Data were processed using NMRPipe^29^ and analyzed with CCPNmr^30^. For residue assignment, a sample of 384 µM ^15^N^13^C-labeled His_6_-mouse Aire CTT in NMR Sample Buffer supplemented with 5% D_2_O was transferred into a 5 mm Shigemi tube. A 2D 15N-HSQC and a set of triple resonance assignment experiments (HNCA, HncoCA, HNCO, HncaCO, HNCACB, hCCCcoNH) were recorded. ^15^N^13^C-labeled His_6_-mouse Aire CTT from the same preparation was mixed with unlabeled CBP TAZ2 and the complex was dialyzed in NMR Sample Buffer. A 2D 15N-HSQC and the same set of triple resonance assignment experiments mentioned above were run on this Aire:CBP sample (ratio 1:1.75). To assign the region showing exchange broadening, 50 mg urea (~1 M) was added in the ^15^N^13^C-labeled His_6_-mouse Aire CTT. The experiments were repeated, leading to 98% assignment of non-proline (non-His_6-_tag) residues. These assignments were then transferred to unbound His_6_-mouse Aire CTT, resulting in assignment of 81% of non-proline residues.

***Fluorescence polarization peptide binding assay***

N-terminal 5,6-carboxyfluorescein (5,6-FAM) FAM-labeled Histone H3 peptide (aa 1-21) with either no K4 methylation or K4me1 were purchased from Anaspec. Binding reaction mixtures (150 μL) contained 50 nM fluorescein-labeled peptide and increasing concentrations of PHDs (0–900 μM) in SEC Buffer. Fluorescence polarization for each sample was measured at 25 °C using a Synergy H1 (BioTek). Data were fit to a simple binding isotherm using Prism (GraphPad Software). Polarization values were normalized to the value without added PHD. All binding experiments were performed at least three times.

***Isothermal titration calorimetry (ITC)***

Protein samples were dialyzed in 20 mM HEPES pH 7.5, 100 mM NaCl, 1 mM DTT for 16 hrs at 4°C. ITC experiments were performed using a VP-ITC calorimeter (MicroCal) and were conducted at 20°C with 19-25 1-2 μL injections, 60 sec delay in between injections with 760μM of mouse Aire CTT in the syringe and 19-76μM TAZ2 in the cell. As a control, 760μM mouse Aire CTT was titrated into the cell containing dialysis buffer only. Thermograms were fit to a two-site binding model using MicroCal Origin 7.0 software. Experiments were collected in duplicate.

***Quantification and statistical analyses***

All statistical analyses for qPCR and fluorescence microscopy quantitation were performed in Prism version 9 or higher (Graphpad). RNA-FISH and IF correlation coefficient was computed in MATLAB (MathWorks). For CRISPR screening, the statistical analysis of sgRNA enrichment was performed using MAGeCK-VISPR (v0.5.6) with the "MAGeCK-RRA" experimental configuration.

Differential expression analysis of RNA-seq data was performed using Rsubread (v2.12.3) and DEseq2 (v1.38.3) R packages. ChIP-seq signals were quantified and normalized using DiffBind (v3.8.4) R package. Statistical analysis details for individual experiments can be found in the figure legends. *p*-value summary: not significant, ns *p*>=0.05; *p* = 0.01-0.05 (*, significant); *p* = 0.001-0.01 (**, very significant); *p* = 0.0001-0.001 (***, extremely significant); *p* <= 0.0001 (****, extremely significant).

***Resource availability***

Further information and requests for resources and reagents should be directed to and will be fulfilled by the lead contact, Sun Hur. All plasmids generated in this study are available from the lead contact with a completed Materials Transfer Agreement.

***Data and code availability***

The accession numbers for the next generation sequencing data reported in this paper is Gene Expression Omnibus: GSE243825. Any additional information required to reanalyze the data reported in this paper is available from the lead contact upon request.

***References for Methods***

11.	Andrews, S. FastQC: A Quality Control Tool for High Throughput Sequence Data [Online]. 2015.

12.	Dobin, A. *et al.* STAR: ultrafast universal RNA-seq aligner. *Bioinformatics* **29**, 15-21 (2013).

13.	Liao, Y., Smyth, G.K. & Shi, W. The R package Rsubread is easier, faster, cheaper and better for alignment and quantification of RNA sequencing reads. *Nucleic acids research* **47**, e47 (2019).

14.	Love, M.I., Huber, W. & Anders, S. Moderated estimation of fold change and dispersion for RNA-seq data with DESeq2. *Genome biology* **15**, 550 (2014).

15.	Wickham, H. *ggplot2: Elegant Graphics for Data Analysis*. Springer-Verlag New York, 2016.
